# PAK3 controls the tangential to radial migration switch of cortical interneurons by coordinating changes in cell shape and polarity

**DOI:** 10.1101/2020.07.06.168179

**Authors:** Lucie Viou, Pierre Launay, Véronique Rousseau, Justine Masson, Clarisse Pace, Robert S. Adelstein, X. Ma, Zhengping Jia, Fujio Murakami, Jean-Vianney Barnier, Christine Métin

**Affiliations:** INSERM U1270, Sorbonne University UMR-S 1270, Paris, France; Université Paris-Saclay, CNRS, Neuroscience Paris-Saclay Institute (Neuro-PSI), 91190 Gif-sur-Yvette; Graduate School of Frontier Biosciences, Osaka University, Yamadaoka 1-3, Suita, Osaka 565-0871, Japan; Laboratory of Molecular Cardiology, NHLBI, Bethesda, MD 20814, USA; The Key Laboratory of Developmental Genes and Human Disease, 2 Sipailou Road, Nanjing, China; Neurosciences & Mental Health, The Hospital for Sick Children, 555 University Ave, Toronto, Canada

**Keywords:** mouse embryonic cortex, organotypic slice, co-culture, videomicroscopy, in utero electroporation, p21-activated kinase, cytoskeleton, myosin 2B, leading process branching, nucleokinesis, directed migration

## Abstract

During the embryonic development, cortical interneurons migrate a long distance tangentially and then re-orient radially to settle in the cortical plate where they contribute to cortical circuits. Migrating interneurons express PAK3, a p21-activated kinase that switches between active and inactive states and controls interneuron migration by unknown mechanism(s). Here we examined the role of the kinase activity of PAK3 to regulate the migration of cortical interneurons. We showed that interneurons expressing a constitutively active PAK3 mutant (PAK3-ca) oriented preferentially radially in the cortex, extended short leading processes and exhibited unstable polarity. On the contrary, interneurons expressing an inactive PAK3 mutant (PAK3-kd for kinase dead) extended branched leading processes, showed directed nuclear movements and remained in the tangential pathways. Results showed that PAK3 kinase activity controls the switch between the tangential and radial modes of migration of cortical interneurons and identified myosin 2B as an effector of this switch.

## INTRODUCTION

The activity of inhibitory GABAergic interneurons (INs) controls the firing of excitatory neurons in cortical circuits. Cortical INs are born in the medial and caudal ganglionic eminences (MGE, CGE) and in the pre-optic area (POA) of the basal telencephalon (Review in Wonders and Anderson, 2006). They reach the cortical plate after a long and complex journey in the embryonic forebrain (Marín and Rubenstein, 2003; Yozu et al., 2005). Within the embryonic cortex, INs first undergo a phase of tangential migration, either at the brain surface in the marginal zone (mz) or deeper within the intermediate and subventricular zones (IZ/SVZ) under the control of extrinsic cues, physical and chemical, either diffusible or contact-dependent (Marín et al., 2010). To enter the cortical plate where they settle and differentiate, INs leave their tangential routes by operating a tangential to radial switch and thereafter migrate radially within the cortex (Faux et al., 2012; Tanaka et al., 2003) (Fig. 1-A). The tangential to radial migratory switch of cortical interneurons is a trajectory re-orientation operated in response to extrinsic guidance cues by highly coordinated cytoskeletal and adhesion reorganizations and by changes in cell motility and cell polarity (Lysko et al., 2011; Martini et al., 2009; Peyre et al., 2015).

**Figure 1:**
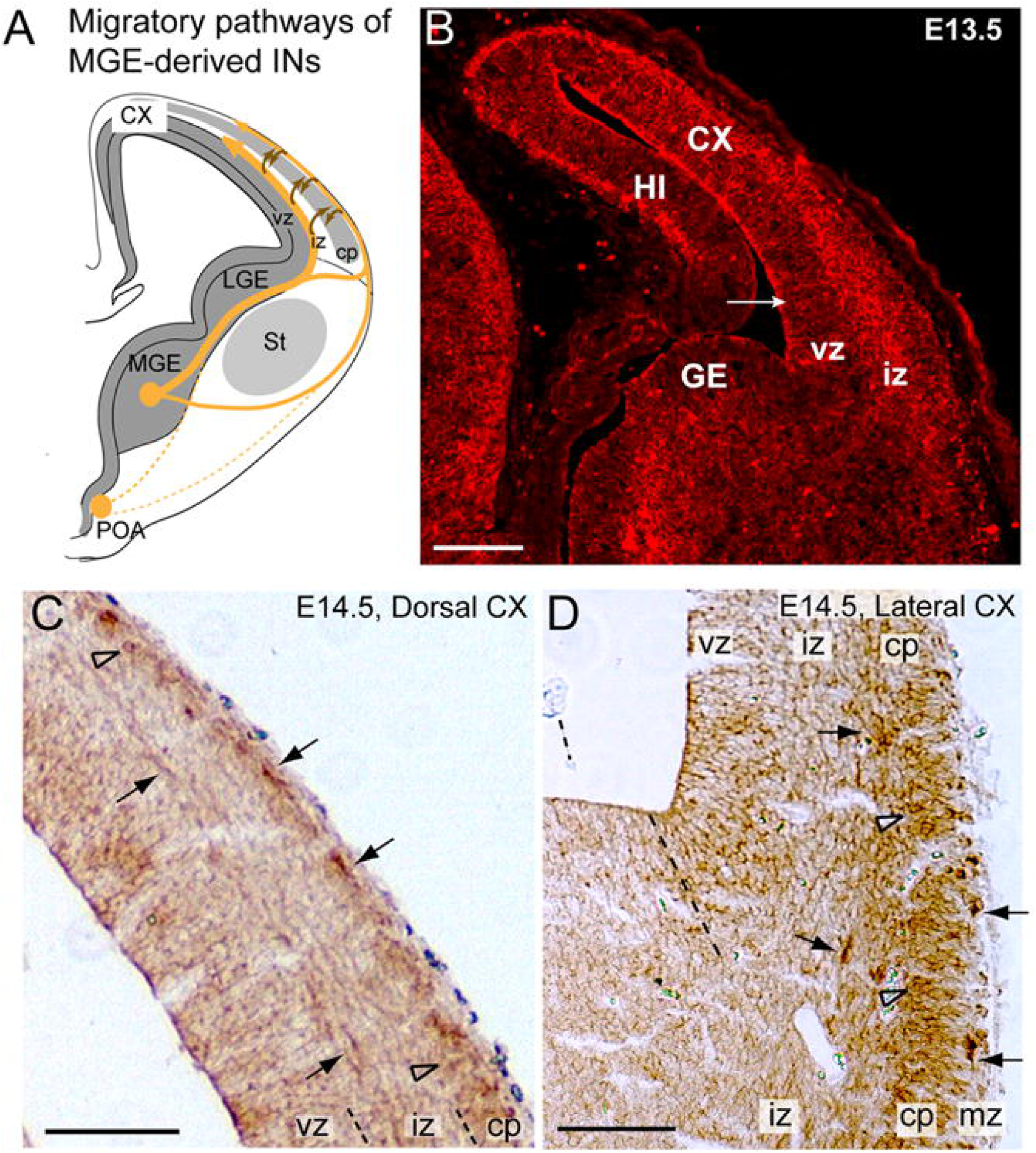
PAK3 expression in the developing cortex of mice embryos at stages E13.5 (B) and E14.5 (C, D). **A.** Scheme illustrating the tangential pathways (yellow) of INs as they migrate in the developing telencephalon and the radial re-orientation (brown) of IN trajectories towards the cp. **B**. PAK3 immunofluorescence on a frontal section of an E13.5 forebrain. PAK3(+) cells are numerous in the sub-ventricular zone of the GE, in the iz of the developing CX, at the pial surface of the HI, and on the ventricular surface of the CX (arrow). C, D. Enlarged views of the dorsal (C) and lateral (D) CX of an E14.5 wild type frontal section. PAK3 immunopositive(+) revealed by DAB staining processes are tangentially oriented in the dorsal cp, in the mz and iz (C,D, black arrows). Radially oriented PAK3(+) processes are observed in the thickness of the cp (C,D, opened arrow heads), and in the vz. CX, cortex; HI, hippocampus; GE, ganglionic eminence; St, striatum; PAO, preoptic area; cp, cortical plate; mz, marginal zone; iz, intermediate zone; vz, ventricular zone. Scale bars: (B,C,D) 200 μm. See also supplementary Fig. S1

Dynamic changes in microtubule and acto-myosin cytoskeleton and in adhesive interactions with the microenvironment are controlled by signaling pathways in which Rho-GTPases play key roles. In the murine developing cortex, CDC42 and RAC1 regulate the production of polarized protrusions in both progenitors and migrating neurons (Konno et al., 2005; Tivodar et al., 2015; Vidaki et al., 2012). By controlling the centrosome position, CDC42 moreover plays an essential role in controlling the polarity of proliferating, migrating and differentiating neurons (Govek et al., 2018; Konno et al., 2005). As expected, *Rac1* and *Cdc42* loss of function in animal models are responsible for major defaults of telencephalon morphogenesis (Chen et al., 2006; 2009) and human mutations in *RAC1* or *CDC42* genes have been associated with a broad spectrum of morphological abnormalities (Martinelli et al., 2018; Reijnders et al., 2017). Among RAC and CDC42 effectors, the p21-activated kinases PAKs play a central role in a large number of signaling cascades that regulate the adhesion, motility and morphology of cells (Bokoch, 2003; Kreis and Barnier, 2009). PAKs therefore fulfill a crucial regulatory function during neurite growth and cell migration. In the developing mouse cortex, PAK1 has been shown to regulate the polarization, morphology and motility of principal excitatory neurons as they migrate along the radial glia (Causeret et al., 2009). PAK3 expression is restricted to neural cells (Kreis and Barnier, 2009). In the mouse basal forebrain, cortical INs express PAK3 only after they entered the developing cortex owing to *Pak3* repression by DLX1/2. PAK3 overexpression observed in *Dlx1/2* KO mice negatively regulates the tangential migration of cortical INs and promotes neurite growth in these neurons (Cobos et al., 2007; Dai et al., 2014). Consistent with their crucial role in brain development, mutations of *PAK1* and *PAK3* in human patients are associated with intellectual disability (ID), severe epilepsy and with a large range of structural abnormalities including microcephaly, macrocephaly and corpus callosum agenesis (Harms et al., 2018; Horn et al., 2019; Ohori et al., 2020). A large majority of *PAK3* mutations identified in patients target the kinase domain of the protein (Duarte et al., 2020; Qian et al., 2020).

We thus examined how the kinase activity of PAK3 controls the migration of cortical INs. As a consequence of the cycling activity of RAC1 or CDC42, PAK proteins similarly switch between inactive and activated conformations in physiological conditions (Bokoch, 2003). We therefore examined the migratory behavior of INs that over-expressed PAK3 mutants locked either in their active (constitutively active, PAK3-ca) or inactive (kinase dead, PAK3-kd) state. We first characterized the cortical trajectories of INs expressing PAK3 mutants, both *in vivo* using in utero electroporation and *ex vivo*, in grafted organotypic forebrain slices.

In a co-culture model designed to analyze the cellular properties of migrating cortical INs, we analyzed the morphology, nuclear and centrosomal movements and growth cone transformations of transfected INs. Our results show that the expression of PAK3-ca and PAK3-kd mutants altered the morphology and motility of INs and influenced their cortical trajectories in specific and opposite ways, whereas PAK3 ablation (*Pak3* KO) and PAK3-wt overexpression have no effect.

INs expressing the non-activable PAK3-kd were unable to re-orient their trajectories away from tangential paths, whereas INs locked in the active state by the expression of PAK3-ca re-oriented radially and no longer switched back to a tangential mode of migration. Interestingly, we identified Myosin 2B as an important but indirect effector of PAK3 in this process. We thus propose that PAK3 kinase activity favors the switch from the tangential to the radial mode of migration by regulating the morphology and polarity of INs.

## RESULTS

### Interneurons expressing PAK3 mutants exhibit abnormal distributions in the embryonic cortex

PAK3 is expressed early during development in the mouse forebrain, as soon as embryonic day 10.5 (E10.5) (Cobos et al., 2007). On E13.5 forebrain sections, immunopositive PAK3 (PAK3+) cells were observed in the intermediate zone (iz) of the cortex and formed a continuum with PAK3+ cells present in the ganglionic eminence (GE) of the subpallium (Fig. 1-B). Pak3 transcripts appeared enriched in the subventricular zone of the lateral (lge) and caudal (cge) GEs, and in the mantle zone of all GEs (Fig. S1). PAK3+ cell bodies and tangentially oriented processes in the iz and marginal zone (mz) of the cortex identified migrating INs (Fig. 1-C,D).

To investigate the influence of the kinase activity of PAK3 on the migration of GE-derived INs, we inserted the murine PAK3 coding sequence fused to either eGFP or mRFP in a pCAG expression vector. Point mutations were introduced in the kinase domain of the wild type protein (PAK3-wt) to either generate a constitutively active kinase (« ca », T421E) or to abrogate the kinase activity (kinase dead « kd », K297L) (Rousseau, 2002) (Fig. 2-A, see Methods). Hereafter, PAK3 constructs fused to eGFP will be referred to as PAK3-wt, PAK3-ca, PAK3-kd.

**Figure 2:**
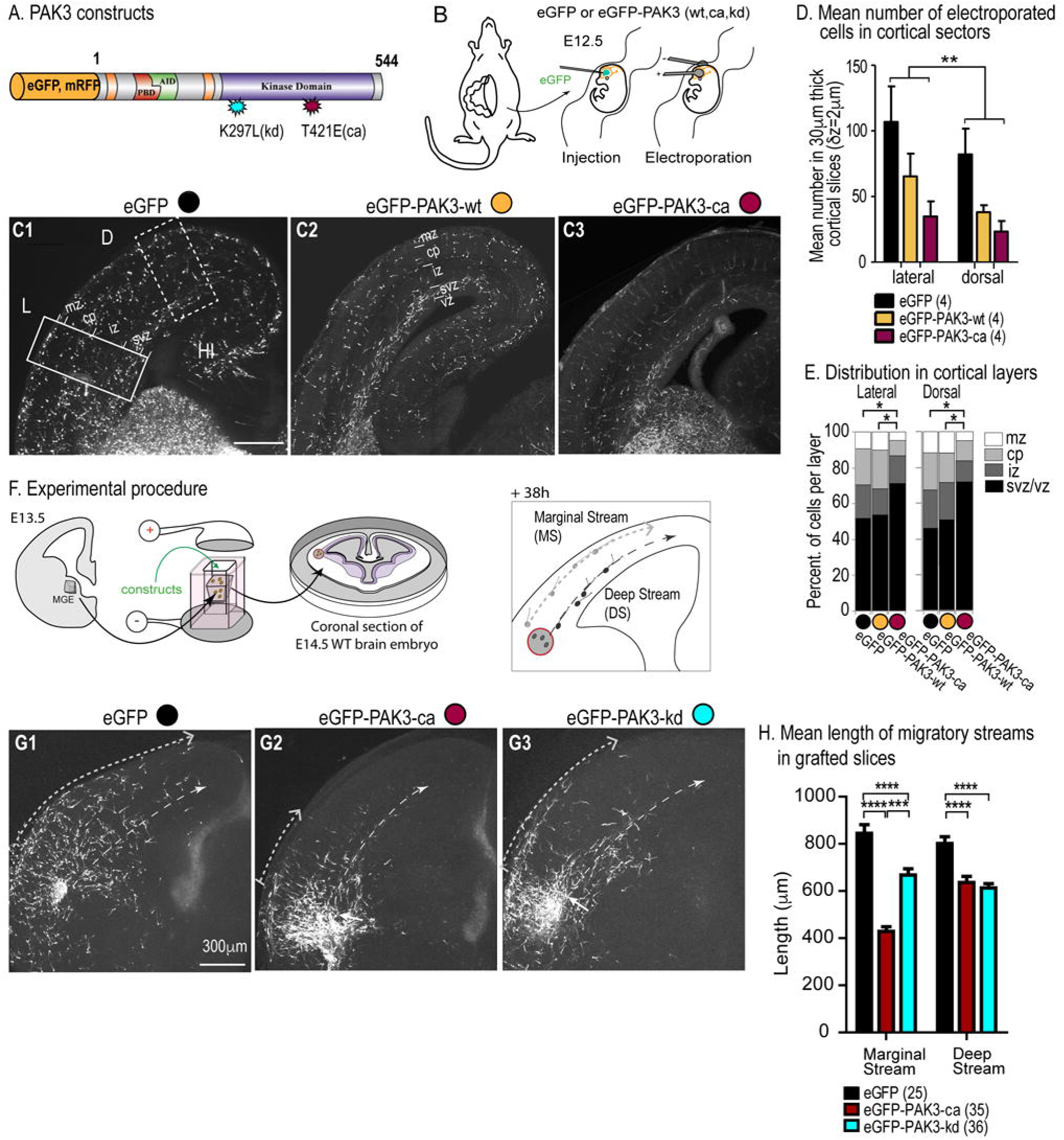
Cortical distribution of interneurons expressing PAK3 mutants in electroporated embryos and in organotypic slices grafted with electroporated MGE explants. **A.** Scheme of the electroporated murine PAK3 constructs. An eGFP or mRFP sequence is fused either to the wild type PAK3 (PAK-wt) or to PAK3 mutants exhibiting a constitutive kinase activity (point mutation T421E, PAK3-ca) or lacking a kinase activity (point mutation K297L, PAK3-kd). **B.** Plasmids were injected in the lower part of the lateral ventricles of E12.5 embryos before electroporating one of the two GE (see Methods). **C1, C2, C3**. Photomicrographs of cortices of fixed E16.5 embryos electroporated at E12.5 in the ipsilateral basal forebrain with an eGFP (C1), eGFP-PAK3-wt (C2) or eGFP-PAK3-ca (C3) construct. Scale bar, 300 μm. **D.** Histograms show the mean number of eGFP+ cells in 300 µm-width lateral (L, solid line rectangle) and dorsal (D, dotted line rectangle) sectors, positioned in the cortex of electroporated embryos as illustrated in C1. Mean number of eGFP+ GE cells significantly differed in the lateral and cortical sectors (Two-way ANOVA, region effect, *F*(1,9)=10.88, p=0.0093). Statistical analysis shows that the number of GE cells electroporated with PAK3 constructs and eGFP plasmid significantly differed (Two-way ANOVA, electroporation effect, *F*(2,9)=4.5, p=0.0442). **E.** Histograms show the distribution of GE cells expressing the different constructs within cortical layers (mz, cp, iz, svz/vz). The distributions of GE cells expressing eGFP and eGFP-PAK3-wt do not significantly differ (Chi2 test with Bonferroni correction, p=1.3068 in lateral cortex, p=1.775 in dorsal cortex). By contrast, the distributions of GE cells expressing eGFP-PAK3-ca and eGFP (eGFP-PAK-wt, respectively) significantly differ in both lateral (p=0.01126, resp. p=0.01425) and dorsal (p=0.0137, resp. p=0.0444) cortex. **F.** Scheme illustrating the experimental procedure to graft an electroporated MGE explant at the pallium/subpallium boundary in a wild type organotypic forebrain slice. After 38h in culture, electroporated MGE cells released from the graft organize two main tangential streams (scheme on the right) i) at the pial surface - marginal stream (MS), light grey dotted line and arrow- and ii) along the ventricle -deep stream (DS), black dotted line and arrow. **G1, G2, G3.** Representative images of cortical slices grafted with MGE explants electroporated with eGFP (G1), eGFP-PAK3-ca (G2) or eGFP-PAK3-kd (G3) constructs. Arrowheads indicate the maximum distance reached by electroporated MGE cells in the marginal and deep streams. **H.** Histograms indicate the mean distance between the most distant electroporated MGE cell and the grafted explant, in the marginal and deep streams. Statistical analysis shows significantly different distances between MGE cells expressing eGFP and either PAK3 mutant (Two-way ANOVA; interaction migratory stream x electroporated construct, *F*(2,93)=32.42; p<0.0001; Bonferroni post-hoc test, p<0.0001). In the marginal stream, distances significantly differ between MGE cells expressing eGFP-PAK3-ca and eGFP-PAK3-kd (Bonferroni post-hoc test, p<0.001). cp, cortical plate; mz, marginal zone; iz, intermediate zone; svz, subventricular zone; vz, ventricular zone. See also supplementary Fig. S2.

Plasmids were electroporated at E12.5 in the GEs of mice embryos as explained in (Luccardini et al., 2013), and the cortical distribution of transfected GE cells was analyzed after fixation on E16.5 brain sections (see Methods, Fig. 2-B,C1-3). As expected, electroporated GE cells that entered the cortex from its lateral edge were significantly more numerous in the lateral than in the dorsal sector (Fig. 2-D). In the cortex of brains electroporated with the PAK3 constructs, either PAK3-wt or PAK3-ca, electroporated cells were less numerous than in the cortex of brains electroporated with a control eGFP plasmid (Fig. 2-D). Nevertheless, the distribution in cortical layers of GE cells electroporated with eGFP and PAK3-wt were remarkably similar (Fig. 2-E). They formed two tangential streams, a large one in the subventricular zone (svz) and a thinner stream in the marginal zone (mz) and moreover distributed in the cortical plate (cp) and in the upper part of the intermediate zone (iz) (Fig. 2-C1,C2). On the contrary, the expression of PAK3 mutants in GE-derived cells strongly altered their cortical distribution (Fig. 2-C3,E, Fig. S2-A). Although the percentage of surviving embryos did not differ in litters electroporated with control (eGFP, PAK3-wt) and mutant (PAK3-ca, PAK3-kd) constructs, PAK3-ca expressing cells were less abundant than PAK3-wt expressing cells. And GE cells electroporated with the PAK3-kd mutant were rarely observed in the cortex at E16.5 (2 out of 30 embryos, Table 1). This result suggests that the expression of inactive PAK3 at an early embryonic stage in GE cells prevented them from crossing the pallium/subpallium boundary to colonize the dorsal pallium *in vivo*. Interestingly, we noticed that the cortical distribution of GE cells with *Pak3* ablation in *Pak3* KO embryos did not differ from the cortical distribution of control GE cells in wild type embryos, at the same stage (Table S1, Fig. S2-B,C,D).

**Table 1:** Number of embryos electroporated in the ganglionic eminences (GEs) at E12.5 with the different constructs. Proportion of embryos that survived until E16.5 and in which electroporated GE cells reached the cortex. **See also supplementary Table S1**

| Plasmids | Number of electroporated embryos | Nber of surviving embryos | Nber of surviving embryos with cortical streams of electroporated GE cells | Mean Nber of cells / embryo (lateral + dorsal sectors) |
| --- | --- | --- | --- | --- |
| pCAG-eGFP | 32 (3 litters) | **27** (84%) | 12 (44% of surviving) | 195 ± 45 |
| pCAG-eGFP-PAK3-wt | 34 (4 litters) | **17** (50%) | 6 (35% of surviving) | 103 ± 22 |
| pCAG-eGFP-PAK3-ca | 51 (5 litters) | **28** (55%) | 7 (25% of surviving) | 58 ± 18 |
| pCAG-eGFP-PAK3-kd | 39 (5 litters) | **30** (77%) | 2 (7% of surviving) |  |

To investigate how PAK3 mutations affect the migration of INs within the cortex, we adopted an ex vivo approach that enables to bypass the subpallium-pallium frontier. We grafted electroporated MGE explants at the pallium-subpallium boundary or in the lateral cortex of organotypic forebrain slices (Fig. 2-F). Electroporated cells released from grafted MGE explants colonized the host cortex. MGE cells expressing the eGFP control construct formed a marginal stream at the surface of the slice in the presumptive mz and a deep stream along the ventricular surface, in the presumptive vz/svz. Numerous individual cells moreover distributed in-between (Fig. 2-G1). Although MGE cells electroporated with either PAK3 mutants migrated in the host cortex, their migration distance was significantly reduced in both marginal and deep streams, as compared with that of control cells (Fig. 2-G2,G3,H). In addition, MGE cells expressing PAK3-ca migrated significantly shorter distances in the marginal stream than MGE cells expressing PAK3-kd (Fig. 2-H). Numerous cells expressing the PAK3-ca mutant distributed in the cp but did not spread tangentially in the mz (Fig. 2-G2), suggesting either that they did not migrate efficiently in the tangential direction or did not reorient tangentially. On the contrary, MGE cells expressing the PAK3-kd construct spread in the marginal and deep streams but poorly colonized the cp located in between (Fig. 2-G3), suggesting that these cells hardly re-oriented radially and persisted in the tangential direction.

### MGE-derived cells expressing different PAK3 mutants follow distinct trajectories in organotypic cortical slices

To confirm these hypotheses, we made a dynamic analysis of the migratory behavior of electroporated MGE cells in organotypic forebrain slices using time-lapse videomicroscopy. We analyzed the trajectories of grafted MGE cells, the dynamics of polarity reversals, and the leading process orientation in fixed cortical slices. Control MGE cells expressing eGFP migrated towards the medial cortex away from MGE explants located at the pallium-subpallium boundary or in the lateral cortex (Fig. 3-A1). They exhibited three main categories of trajectories: numerous cells migrated tangentially in the deeper part of the cortex (vz/svz, red trajectories, Fig. 3-B1), a few cells migrated tangentially in the mz (pink trajectories, Fig. 3-B1) and another large group of cells migrated first tangentially in the deep part of the cortex and then reoriented either radially or with an angle to the cp (green trajectories in Fig. 3-B1).

**Figure 3:**
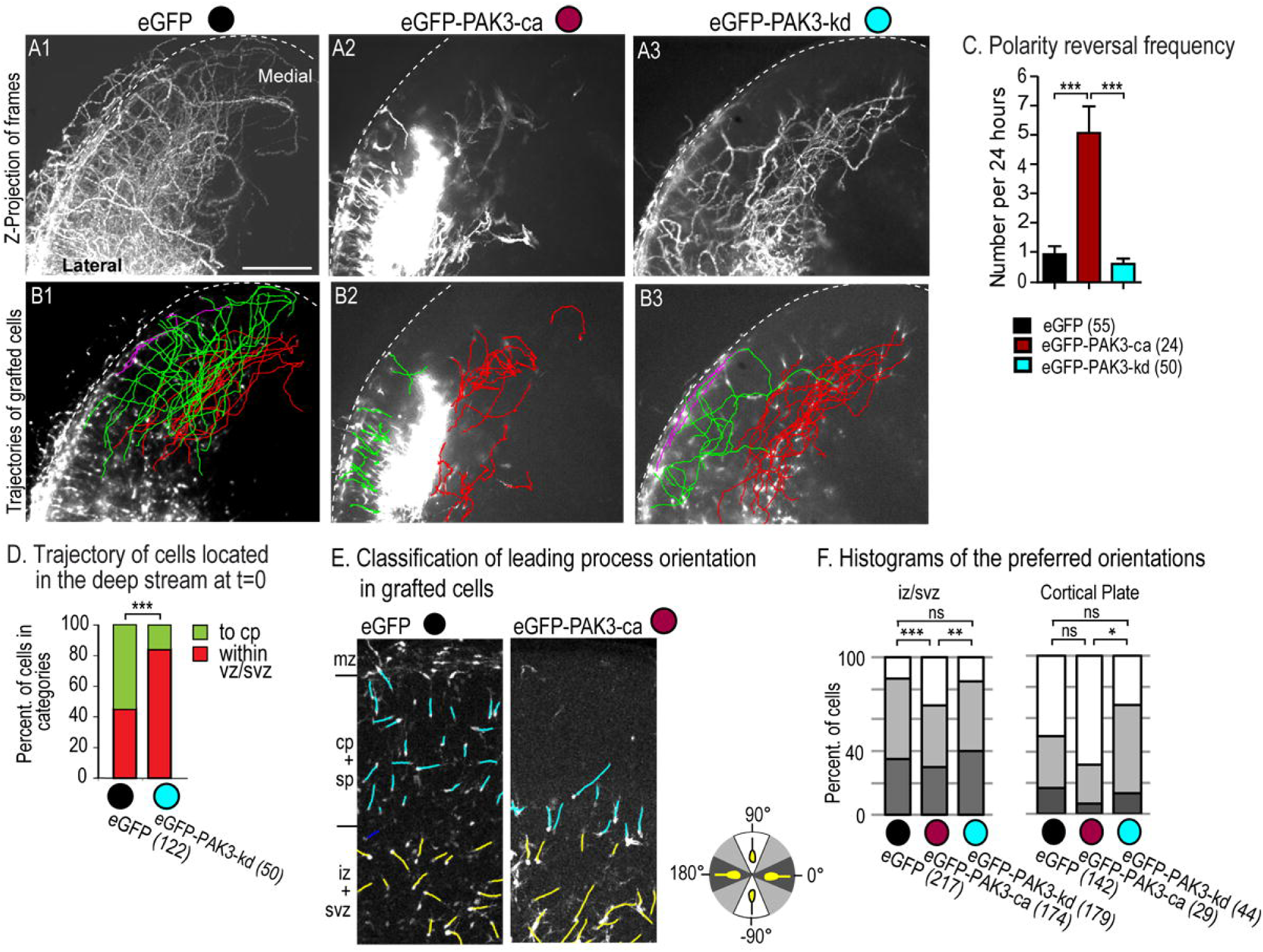
Trajectories of MGE cells electroporated with PAK3 mutants in cortical slices A1, A2, A3, B1, B2, B3. MGE cells electroporated with eGFP (A1, B1), eGFP-PAK3-ca (A2, B2) or eGFP-PAK3-kd (A3, B3) constructs and migrating in a host cortical slice were imaged every 10 minutes for 24 hours. **A1-3** are z-projections of recorded stacks. See also supplementary Movie3A1, Movie3A2, Movie3A3. **B1-3** show color-coded manually tracked trajectories of cells in a same slice: red trajectories remain below the cortical plate (cp) in the deep stream, green trajectories are directed from the deep stream to the cp, and pink trajectories keep above the cp, in the superficial stream. Scale bar (A1), 300 μm **C.** Histogram shows that polarity reversals (inversion of nuclear movement direction) were significantly more frequent in MGE cells expressing eGFP-PAK3-ca than in MGE cells expressing eGFP and eGFP-PAK3-kd (One-way ANOVA, Dunn’s test, p<0.001). **D.** Histogram compares the frequency of trajectories directed to the cp (green) or confined within the vz/svz (red) in MGE cells expressing eGFP and eGFP-PAK3-kd (Fisher test, p<0.001). **E, F.** Leading process orientation in fixed cortical slices of grafted MGE cells expressing eGFP and PAK3 constructs. **E.** Leading process orientation was measured with regards to the cortical surface and angles were classified as radial (90°± 22.5°, white sector), oblique (45°±22.5°, grey sector) or tangential (0°±22.5°, black sector) as illustrated in scheme. **F.** Histograms show the frequency of each class of orientation in the iz/svz (left histogram) and cp (right histogram). In the iz/svz, orientations significantly differed between MGE cells expressing eGFP-PAK3-ca and either eGFP (Chi2 test, p=0.0009) or eGFP-PAK3-kd (Chi2 test, p=0.0038). In the cp, distributions significantly differed in MGE cells expressing eGFP-PAK3-kd and eGFP-PAK3-ca (Fisher test, p= 0.0251). See also supplementary Fig. S3.

By contrast, MGE cells expressing PAK3-ca did not migrate long distances away from their explant of origin (Fig. 3-A2). Some cells migrated radially in the cp (green trajectories, Fig. 3-B2). A minimal population moved in the lower iz/svz but displayed short and curved trajectories (red trajectories, Fig. 3-B2). Polarity reversals (inversion of nuclear movement direction) were 5 times more frequent than in control cells (Fig. 3-C, red bar). Interestingly, the large majority of MGE cells expressing PAK3-kd migrated tangentially in the lower iz/svz (Fig. 3-A3,B3). These cells moved significantly less frequently to the cp than MGE cells expressing control eGFP (Fig. 3-D, blue dot). The trajectories of MGE cells expressing PAK3-kd appeared slightly less straight than those of control eGFP cells (Fig. 3-B3).

In agreement with these observations, PAK3-ca expressing MGE cells grafted in cortical slices and cultured for 38 hours before fixation possessed leading processes that were more frequently radially-oriented in the cp (90°±22.5°) than MGE cells expressing eGFP or PAK3-kd (Fig. 3-E,F, right hand histogram). In the iz/svz, leading processes of PAK3-ca expressing MGE cells where less frequently oriented tangentially or with an angle (0°±22.5° or 45±22.5°) than those of MGE cells expressing eGFP or PAK3-kd (Fig. 3-E,F left hand histogram).

In summary, MGE cells expressing PAK3-ca frequently reversed their polarity and showed an abnormal tangential migration in the iz/svz. By contrast, MGE cells expressing PAK3-kd migrated long distances tangentially in the deep iz/svz of the host slice but showed an altered ability to colonize the cp. As expected, MGE cells expressing PAK3-ca were more frequently radially-oriented in the cortical plate than either eGFP or PAK3-kd expressing MGE cells (Fig. 3-F).

### The trajectories of electroporated MGE cells in co-cultures share features with in vivo trajectories

The above differences could reflect differential responses of MGE cells expressing different PAK3 mutants to the three-dimensional structure of the host cortex. Alternatively, they could identify intrinsic differences in the migratory properties of transfected MGE cells that will be conserved in flat co-cultures. To distinguish between these two hypotheses, we analyzed the migratory behavior of electroporated MGE cells in a flat co-culture model developed and currently used in the laboratory (Bellion et al., 2005) (Fig. 4-A, Fig. S4-A). Electroporated MGE explants were cultured on dissociated wild type cortical cells plated on laminin-coated glass coverslips. Released MGE cells migrated centrifugally away from their origin explant. Here, we imaged migrating MGE cells by time-lapse videomicroscopy for an average of 24h and analyzed trajectory properties and cell directionality (Fig. 4-B,C,D, Fig. S4-C,D,E). Most MGE cells expressing the control eGFP construct exhibited straight trajectories (Fig. 4-E, 60% of recorded cells). The proportion of straight, curved and confined trajectories significantly differed between MGE cells expressing eGFP and PAK3 mutants. MGE cells expressing PAK3-ca showed an increased proportion of confined trajectories due to frequent polarity reversals (Fig. 4-E middle bar), whereas MGE cells expressing PAK3-kd showed an increased proportion of curved trajectories (Fig. 4-E right hand bar).

**Figure 4:**
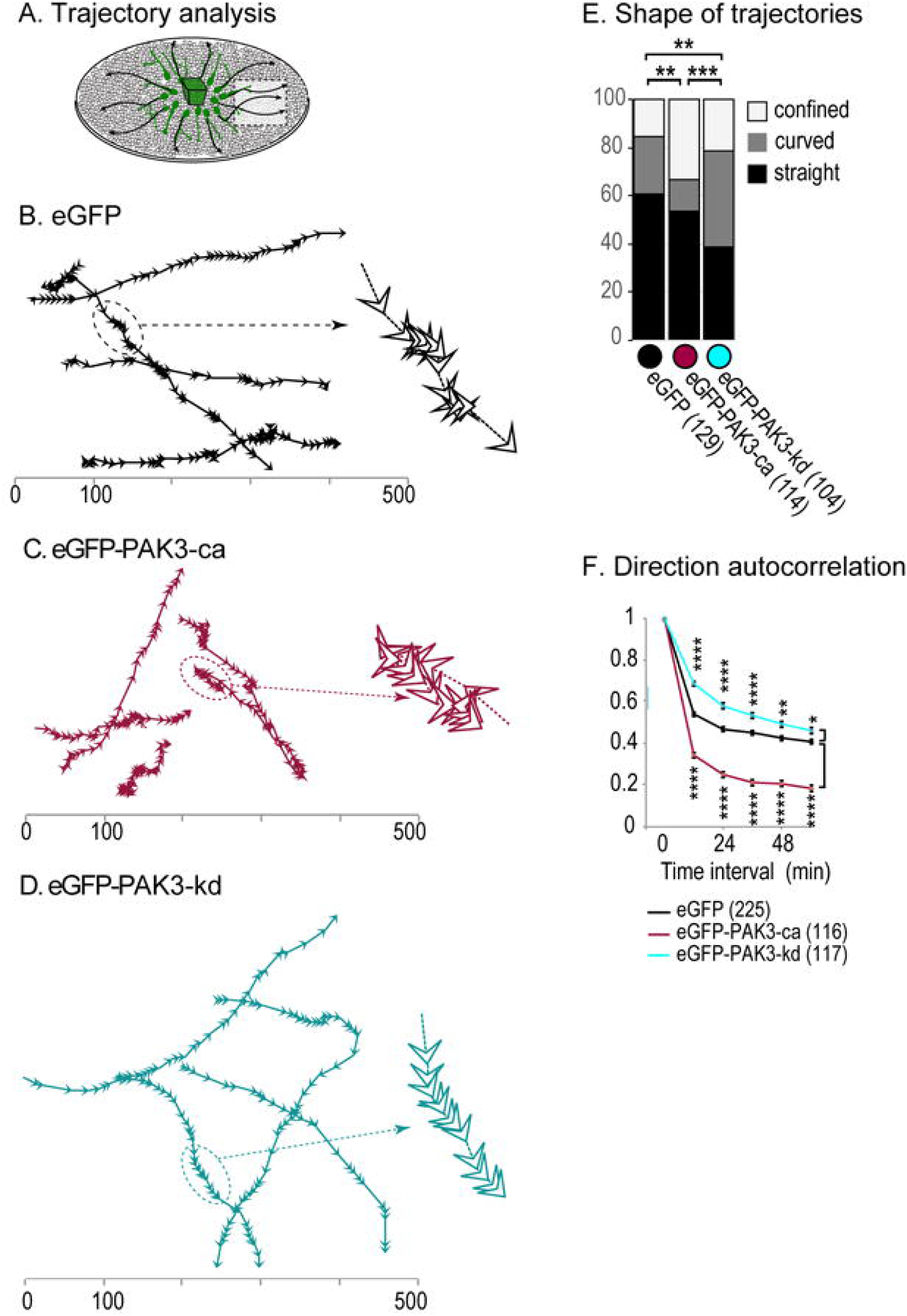
Trajectories of MGE cells electroporated with PAK3 mutants in co-cultures. **A.** Scheme of the co-culture model (see Fig. S4-A). Cells migrated centrifugally away from MGE explants on the substrate of cortical cells. MGE cells were imaged at the migration front (white area) every 3 minutes with a X20 objective for at least 20 hours. **B, C, D.** Representative trajectories of MGE cells expressing eGFP (B), eGFP-PAK3-kd (C) and eGFP-PAK3-ca (D) constructs. Portions of trajectories are enlarged on the right side. Arrowheads indicate cell body position and the direction of movement. See also supplementary Movie4B, Movie4C, Movie4D. **E.** Histogram representing the proportion of straight (black: during the recording session, MGE cells enter and exit the field of view by opposite sides), curved (grey: during the recording session, MGE cells enter and exit the field of view by adjacent sides) and confined (white: during the recording session, MGE cells remain in the field of view) trajectories in MGE cells expressing eGFP (black dot), eGFP-PAK3-ca (red dot) and eGFP-PAK3-kd (blue dot) constructs. Proportions significantly differed between MGE cells expressing eGFP and eGFP-PAK3-ca (Chi2 test, p=0.0077), eGFP and eGFP-PAK3-kd (Chi2 test, p=0.0054), eGFP-PAK3-ca and eGFP-PAK3-kd (Chi2 test, p=0.00015). **F.** Curves illustrate the direction autocorrelation coefficient (cosine of the angular deviation of MGE cell trajectories when the time interval between cell positions increases, Gorelik and Gautreau, 2014) for MGE cells expressing eGFP and eGFP-PAK3 mutant constructs). Coefficient is 1 for straight trajectories. The number of analyzed trajectories is indicated. Statistical analysis by Two-way ANOVA shows an effect of the interaction construct x time interval (*F*(10,2730)=29.32, p<0.0001). Post-hoc Bonferroni tests show significantly different coefficients for time intervals comprised between 12 and 60 minutes. See also supplementary Fig. S4.

We thereafter quantified the cell directionality by analyzing the angular deviation of trajectories expressed as a “direction autocorrelation coefficient” calculated using a macro from (Gorelik and Gautreau, 2014) (Fig. 4-F). This coefficient is close to 1 for straight trajectories. As expected, the coefficient was smaller for PAK3-ca expressing cells that frequently reverse their polarity than for control cells. Moreover, the persistence of direction of MGE cells expressing PAK3-ca decreased much faster with time than that of control MGE cells expressing eGFP (Fig. S4-F). Surprisingly enough, the autocorrelation coefficient of MGE cells expressing PAK3-kd was significantly higher than in control eGFP MGE cells, indicating that PAK3-kd expressing MGE cells were significantly more directional than control eGFP MGE cells (Fig. 4-F). Accordingly, the persistence of direction index of these cells remained above control values (Fig. S4-F). Closer examination of trajectories confirmed that PAK3-kd expressing MGE cells were more processive than control eGFP MGE cells that frequently performed transient and short duration polarity reversals (compare enlarged trajectories in Fig. 4-B,D).

In summary, MGE cells expressing PAK3-ca frequently reversed their polarity and this property restrained their displacements in co-cultures whereas INs expressing PAK3-kd showed long processive and possibly curved trajectories. PAK3 mutants thus similarly affected the migratory properties of MGE cells in co-cultures and organotypic slices.

### The expression of PAK3 mutants alters leading process morphology in MGE cells

Directional changes of INs have been correlated with leading process branching (Lysko et al., 2011; Martini et al., 2009). We thus examined whether the leading process morphology (length, branching pattern) of MGE cells transfected with PAK3 mutants was modified in addition to the trajectory alterations observed in co-cultures and cortical slices. Analyses were performed at the migration front in fixed co-cultures.

MGE cells expressing PAK3-ca had significantly shorter leading processes than control and PAK3-kd expressing MGE cells (Fig. 5-A1-3,B). Regardless of the electroporated construct, MGE cells showed branched leading processes (Fig. 5-A1-3). However, leading processes were significantly more branched in MGE cells expressing PAK3-kd than in control and PAK3-ca expressing MGE cells (Fig. 5-C). Interestingly, in contrast with *Pak3* ablation that did not affect the maximal leading process length, PAK3-wt overexpression moderately but significantly decreased this parameter (Fig. S5-A1,A2). Neither *Pak3* ablation (Fig. S5-B) nor PAK3-wt overexpression (not illustrated) significantly altered the branching pattern of MGE cells.

**Figure 5:**
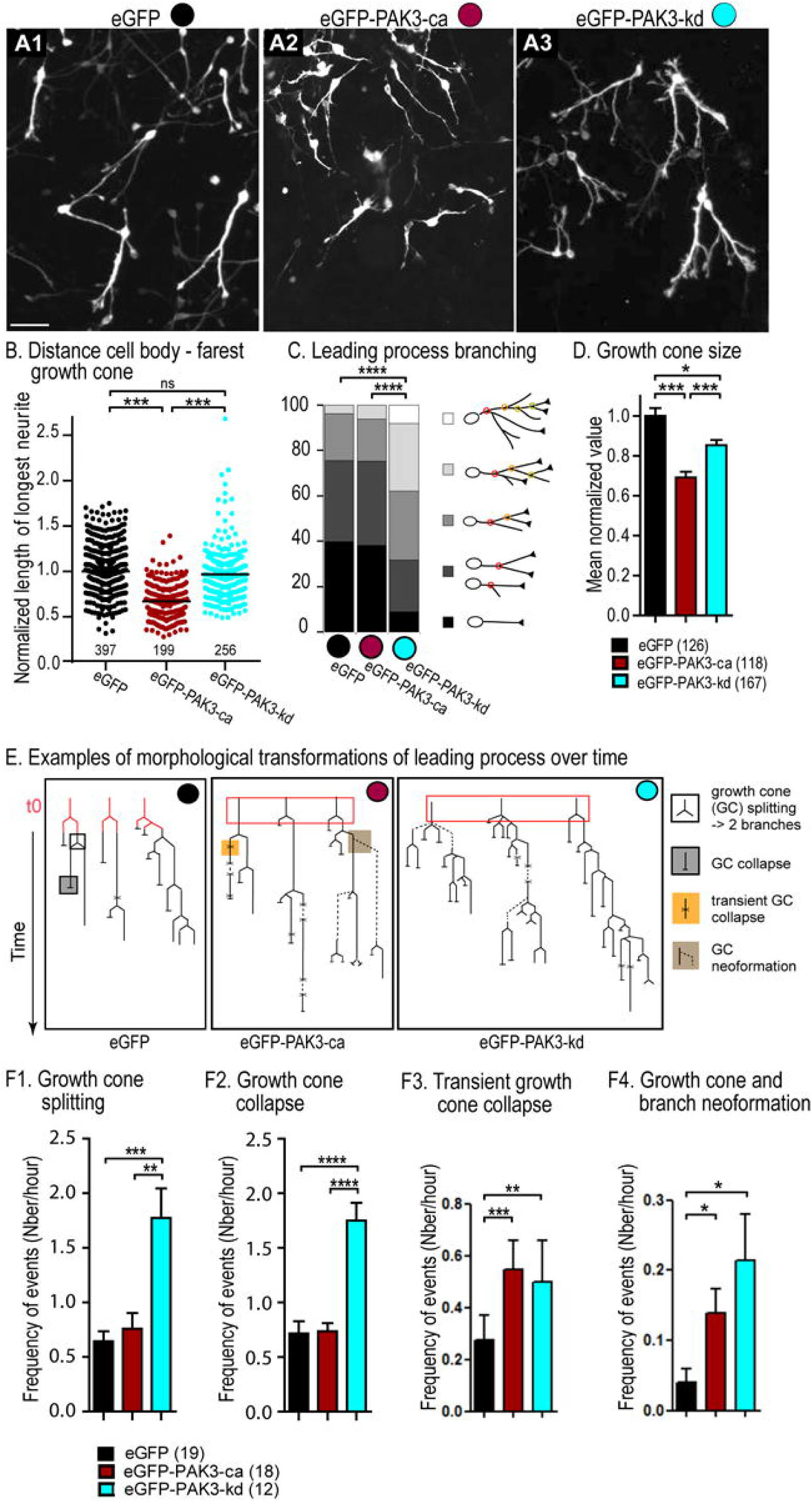
Leading process morphology and growth cone dynamics in MGE cells expressing PAK3 mutants. **A1, A2, A3.** Photomicrographs of MGE cells imaged at the migration front of co-cultures (see Fig. 4-A). MGE cells express control eGFP (A1, black dot), eGFP-PAK3-ca (A2, red dot) and eGFP-PAK3-kd (A3, blue dot) constructs. Scale bar, 20 μm. **B.** Graphs showing the length of the longest neurite in large samples of electroporated cells (number indicated below graphs). Lengths were normalized to the mean value in the control sample (eGFP expressing cells). Statistical significance tested by One-way ANOVA and post-hoc Dunn’s tests. **C.** Histogram showing the percentage of MGE cells exhibiting a leading process without (black), with 1 (dark grey), 2 (medium grey), 3 (light grey), 4 or more (white) bifurcations (Fisher tests for comparisons). **D**. Histogram showing the mean surface (pixel number) of growth cones at their maximum size on movies. Values were normalized with regards to the mean growth cone size in MGE cells expressing eGFP (Kruskal-Wallis test, p<0.001 and post-hoc Dunn’s tests for pair comparisons). **E**. Dendograms schematize the leading process transformations of MGE cells electroporated with eGFP (black dot), eGFP-PAK3-ca (red dot), and eGFP-PAK-kd (blue dot) constructs over time. Four main growth cone transformations indicated in the legend were identified and their frequency calculated. See also supplementary Movie5E. **F1, F2, F3, F4.** Histograms give the mean frequency of leading growth cone splitting (F1), collapse (F2), transient collapse (F3) and growth cone neoformation leading to a novel branch (F4) in MGE cells expressing eGFP (19 cells), eGFP-PAK3-ca (18 cells), and eGFP-PAK3-kd (12 cells) constructs, recorded for more than 2 hours. In F1 and F2, statistical significance is assessed by One-way ANOVA and post-hoc Dunn’s tests. In F3 and F4, statistical significance is assessed by Poisson-ANOVA model, and post-hoc likehood ratio test, *** p=0.0007, ** p=0.0031 in F3, and * p=0.036, p=0.0116 in F4. See also supplementary Fig. S5.

The leading process morphology reflects the activity of growth cones at the tip of neurites. We therefore analyzed growth cone size and dynamic transformations (division, collapse and neoformation) on time-lapse movies of MGE cells expressing eGFP, PAK3-ca and PAK3-kd. We observed that growth cones were significantly smaller in MGE expressing PAK3-ca than in MGE cells expressing eGFP and PAK3-kd (Fig. 5-D). In control MGE cells, the most frequent events were i) growth cone splitting that increased the number of neurites and ii) growth cone collapse that pruned supernumerary neurites (Fig. 5-E,F1,F2, black dot, black bars). Transient collapses followed by neuritic regrowth were much less frequent (Fig. 5-F3, black bar). MGE cells expressing PAK3-kd, which presented highly branched leading processes, exhibited twice as much growth cone splitting than control cells and also more frequent growth cone neoformation (Fig. 5-E,F1,F4, blue dot, blue bars). In these neurons, the frequency of collapses and of transient collapses was increased accordingly (Fig. 5-F2,F3, blue bars). In MGE cells expressing PAK3-ca, whose leading process branching did not differ from control MGE cells, the main change was an increased transient collapse frequency and a slight increase in growth cone neoformation (Fig. 5-E,F1,F2,F3,F4, red dot and red bars).

In summary, PAK3-ca expressing MGE cells that frequently reversed their polarity during migration had shorter leading processes and smaller growth cones than control MGE cells. PAK3-kd expressing MGE cells that migrated long distances tangentially and showed more direction persistence, maintained highly branched leading processes thanks to the balance in growth cone production and elimination.

### The expression of PAK3 mutants alters centrosomal and nuclear movements in MGE cells

In migrating neurons, the leading process exerts a pulling force on the centrosome, whose forward migration controls nuclear progression (Peyre et al., 2015; Solecki et al., 2009) (Fig. 6-A1,A2). We therefore examined centrosomal movements in MGE cells expressing PAK3 mutants, owing to their altered leading process morphology. Dynamic analyses by time-lapse videomicroscopy of MGE cells co-transfected with a fluorescent centrosomal marker (pericentrin-mKO1) revealed that the forward centrosomal movements were significantly shorter in MGE cells co-expressing PAK3-ca and significantly longer in MGE cells co-expressing PAK3-kd as compared to pericentri-mKO1/eGFP co-expressing MGE cells (Fig. 6-B2,B3,C). Accordingly, the cytoplasmic swelling that accompanies forward centrosomal movements formed less frequently in MGE cells expressing PAK3-ca and more frequently in MGE cells expressing PAK3-kd than in control eGFP MGE cells (not illustrated).

**Figure 6:**
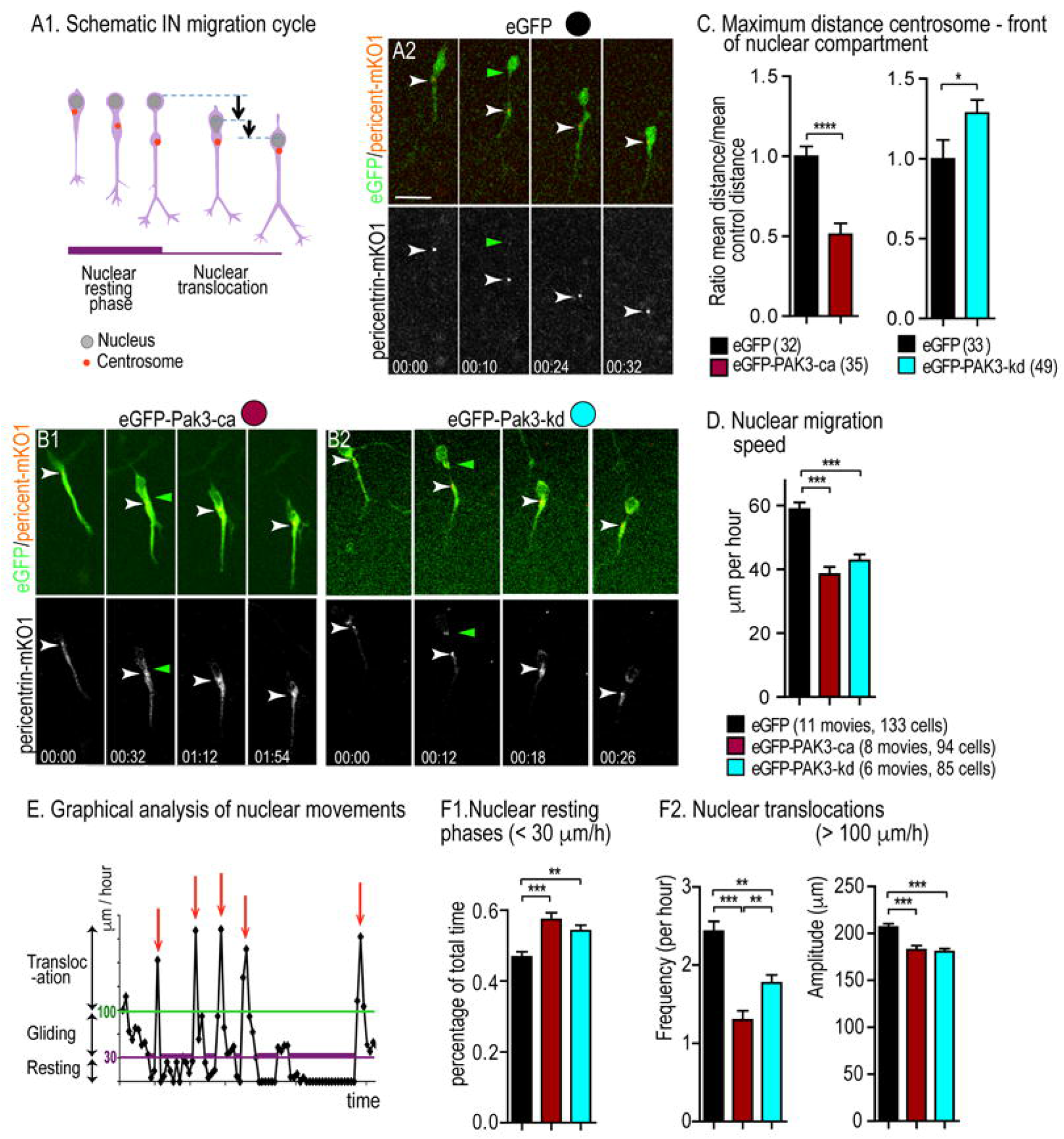
Abnormal centrosomal and nuclear movements in MGE cells expressing PAK3 mutants. **A1, A2.** Centrosomal and nuclear movements in control condition. **A1.** Scheme illustrating two main phases of the migration cycle of cortical INs: 1) the nuclear resting phase during which the centrosome migrates forward in a cytoplasmic swelling, 2) the nucleus translocation towards the centrosome. **A2.** Time-lapse sequence of a migrating MGE cell co-electroporated with the eGFP (green) and pericentrin-mKO1 (red) plasmids. Pericntrin-mKO1 labels the centrosome (white arrowhead). Time is indicated in hours:minutes on frames. The nucleus is the oval structure at cell rear. Scale bar, 10 µm. See also supplementary Movie6A2. **B1, B2.** Time-lapse sequences illustrate the migration of MGE cells co-electroporated with eGFP-PAK3-ca (**B1**), eGFP-PAK3-kd (**B2**) and pericentrin-mKO1 plasmids. See also supplementary Movie6B1 and Movie6B2. **C.** Histograms compare the maximal centrosome-nuclear front distance (between green and white arrowheads in A2, B1, B2) in MGE cells expressing eGFP (black bars), eGFP-PAK3-ca (red bar, left), and eGFP-PAK3-kd (blue bar, right). Statistical significance is assessed by Student T-tests. **D.** Histogram comparing the mean migration speed of the cell body in MGE cells expressing eGFP and PAK3 mutants. Statistical significance is assessed by One way-ANOVA and post-hoc Dunn’s tests. **E.** Graph defining the different modes of nuclear movements in migrating MGE cells. Resting phases of the nucleus are defined by instantaneous speed below 30 microns per hour, gliding movements by instantaneous migration speed between 30 and 100 microns per hour and nuclear translocations (red arrows) by instantaneous speed above 100 micron per hour. **F1, F2.** Histograms compare the percentage of resting phase (F1) and the frequency (F2, left) and amplitude (F2, right) of nuclear translocations in MGE cells expressing eGFP and PAK3 mutants. Same legend as in D. Statistical significance is assessed by One-way ANOVA and post-hoc Dunn’s test. See also supplementary Fig. S6

Despite opposite changes in nuclear–centrosomal distance, MGE cells expressing both PAK3 mutants exhibited decreased nuclear migration speeds (Fig. 6-D). The saltatory progression of nuclei was altered more dramatically in PAK3-ca than in PAK3-kd expressing MGE cells (Fig. 6-E,F1,F2). Nuclear resting phases were much longer and nuclear translocations much less frequent in PAK3-ca than in PAK3-kd expressing MGE cells (Fig. 6-F1,F2 left), but the amplitude of nuclear translocations was similarly affected (Fig. 6-F2 right). MGE cells expressing PAK3 mutants, which were grafted in organotypic cortical slices, presented similar alterations and PAK3-ca expressing MGE cells appeared even more severely affected (Fig. S3).

The mean nuclear migration speed remained unchanged in INs expressing the PAK3-wt construct and in *Pak3* KO INs (Fig. S6-A,H). The only significant defect that we detected was a small but significant decrease of the fast nuclear movement amplitudes (Fig. S6-D,I).

In summary, analyses revealed a tight relationship between the leading process morphology and centrosomal movements: PAK3-ca expressing MGE cells with short leading processes showed short forward centrosomal movements, whereas PAK3-kd expressing MGE cells with developed and branched leading processes showed long forward centrosomal movements. Surprisingly, these opposite changes both associated with an increased nuclear pause duration and a decreased nuclear translocation frequency. Nucleokinesis defects were nevertheless more pronounced in PAK3-ca expressing MGE cells, identifying large forward centrosomal movement amplitudes as a key regulator of the saltatory nucleokinesis.

### The expression of PAK3 mutants alters the activity of Myosin 2B in migrating INs

We previously showed that centrosome mis-positioning is associated with abnormal Myosin 2B dynamics (Luccardini et al., 2013). We therefore analyzed the distribution of Myosin 2B in MGE cells expressing PAK3-ca or PAK3-kd mutants. MGE explants dissected from KI embryos with a GFP sequence inserted in the *Myosin 2B* gene (non-muscle myosin IIB-GFP, Bao et al., 2007) were electroporated with plasmids encoding PAK3 mutants fused to mRFP (hereafter referred to as RFP-PAK3-ca and RFP-PAK3-kd). Control cells were electroporated with mRFP.

Myosin 2B plays a crucial role in tangentially migrating neurons (Ma et al., 2004). Control MGE cells dissected from Myosin 2B-GFP embryos were imaged at rather high frequency (once every minute) in co-cultures to visualize Myosin 2B-GFP dynamics. Analyses revealed stereotyped changes in Myosin 2B-GFP distribution during the migration cycle (Fig. 7-A,B). During the nuclear resting phase when the centrosome moves forward (start of the migratory sequence in Fig. 4-A1), Myosin 2B-GFP speckles distributed around the nucleus and accumulated in the rostral swelling where the Myosin 2B-GFP signal progressively peaked (pattern 1, Fig. 7-B). Prior and during nuclear translocation, the Myosin 2B-GFP signal relocated at the nuclear rear (pattern 2, Fig. 7-B). Following nuclear translocation, the Myosin 2B-GFP shifted from nuclear rear to nuclear front (pattern 3, Fig. 7-B) and thereafter, Myosin 2B-GFP speckles spread in the whole cell body comprising the rostral swelling (pattern 4, Fig. 7-B). Fast nuclear movements were correlated with strong Myosin 2B-GFP signal at the nuclear rear, whereas intermediate nuclear speeds were measured when the Myosin 2B-GFP signal was located at the nuclear front (Fig. S7-A,B,C).

**Figure 7:**
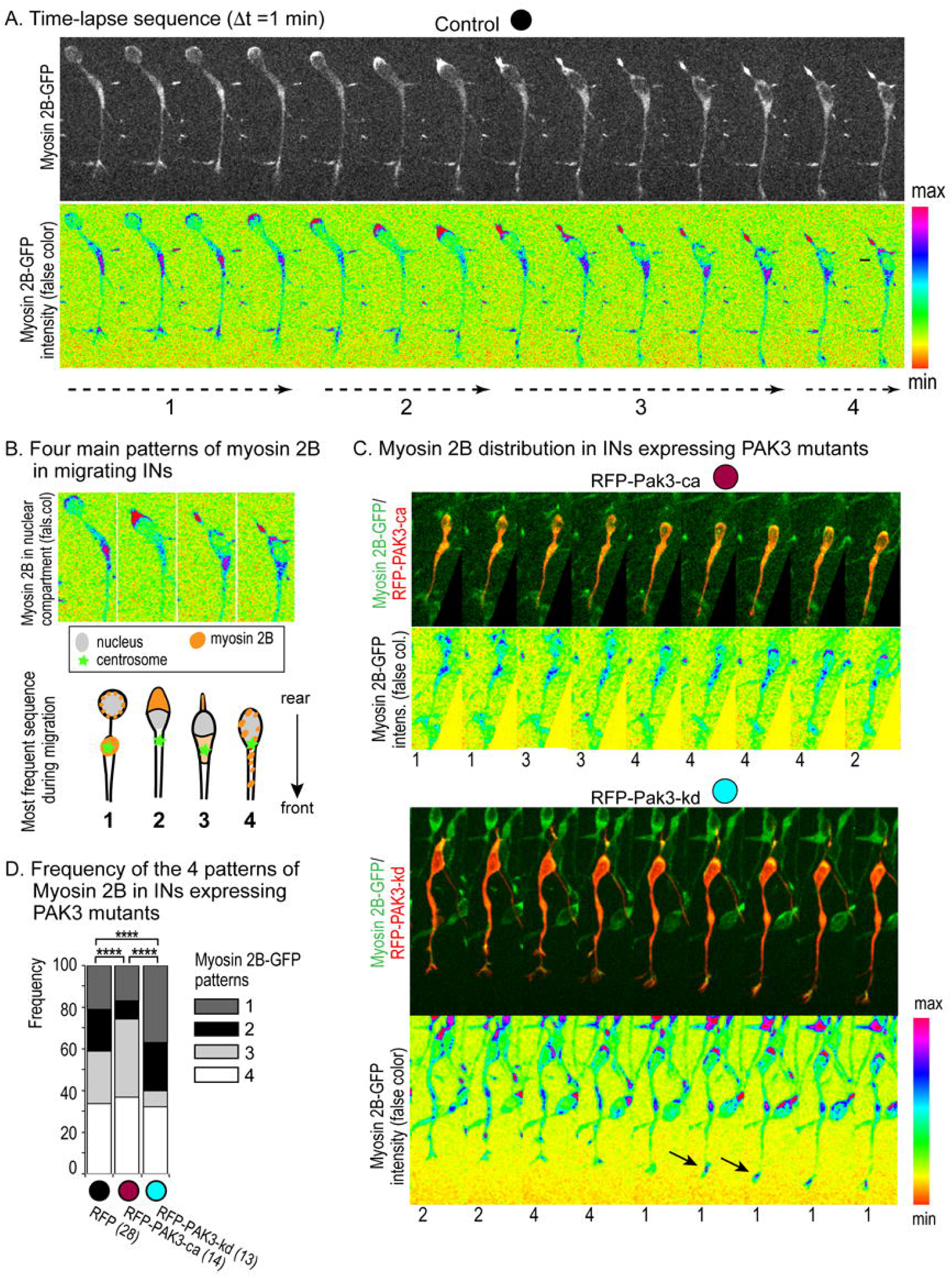
Myosin 2B distribution during the migration cycle in control and PAK3 mutant expressing MGE cells. **A.** Time-lapse sequence of a MGE cell expressing the Myosin 2B-GFP fusion protein migrating in a co-culture (see Fig. 4-A). The bottom row shows the GFP fluorescence intensity color-coded as indicated on scale. See also supplementary Movie7A. **B.** Pictures (top row) and schemes (bottom row) illustrate the most frequent sequence of Myosin 2B distribution in migrating MGE cells (see Fig. S7-D,E): 1, Myosin 2B accumulates in the rostral swelling during centrosomal forward migration (and nuclear resting phase); 2, Myosin 2B relocates at the nuclear rear before and during nuclear translocation; 3, Myosin 2B repositions at the nuclear front after nuclear translocation and during gliding nuclear movement; 4, Myosin 2B speckles spread in the whole cell body during resting phase. See also supplementary Movie7C1 and Movie7C2. **C.** Time-lapse sequences illustrate the distribution of Myosin 2B-GFP in migrating MGE cells electroporated with RFP-PAK3-ca (top) and RFP-PAK3-kd (bottom) constructs. On each frame, Myosin 2B-GFP patterns are numbered according to the sequence described in (B). **D.** Histogram representing the frequency of the four Myosin 2B-GFP patterns in MGE cells expressing RFP (black dot), RFP-PAK3-ca (red dot) and RFP-PAK3-kd (blue dot). Statistical significance of distributions is assessed by Chi2 tests (p<0.0001). See also supplementary Fig. S7

Similar patterns of Myosin 2B-GFP were observed in MGE cells electroporated with RFP-PAK3 mutants (Fig. 7-C). However, the accumulation of Myosin 2B-GFP at the nuclear rear (pattern 2) was observed half as often in MGE cells expressing RFP-PAK3-ca (Fig. 7-C,D, red dot) than in control MGE cells expressing RFP (Fig. 7-D, black dot), in agreement with the decreased frequency of fast nuclear translocations in MGE cells expressing PAK3-ca (Fig. 6-F2 left, red dot). In these cells, Myosin 2B-GFP distributed therefore more frequently at the nuclear front than in control RFP MGE cells (Fig. 7-C,D, red dot, pattern 3). In MGE cells expressing RFP-PAK3-kd, the myosin 2B-GFP signal was more frequently observed in the rostral swelling and at the nuclear rear than in control RFP MGE cells (Fig. 7-C,D, blue dot, patterns 1 and 2). Moreover, a strong but spatially restricted Myosin 2B-GFP signal was often observed in these cells at the tip of neurites during growth cone splitting phases (arrows in Fig. 7-C, blue dot).

In summary, analyses showed that PAK3-ca expression impaired Myosin 2B accumulation at the nuclear rear, whereas PAK3-kd expression maintained Myosin 2B at the nuclear rear, identifying PAK3 as a negative regulator of Myosin 2B concentration at the cell rear in migrating interneurons. PAK3 activity displaced the Myosin 2B signal from the rostral swelling and the nuclear rear where it preferentially positioned in the absence of PAK3 kinase activity.

Altogether, our results suggest that PAK3 activity regulates the terminal distribution of cortical INs by promoting a tangential-to-radial switch of their cortical trajectory through a myosin IIB–dependent mechanism that controls cell morphology, cell polarity and cell dynamics.

## DISCUSSION

### Mutations targeting the kinase domain of PAK3 are responsible for intellectual disability (ID) in humans

PAK3 mutations in humans are responsible for ID, developmental delay including motor and speech deficits, behavioral problems linked to increased aggression and violence, and severe epilepsy (Allen et al., 1998; Horvath et al., 2018; Iida et al., 2019; Magini et al., 2014; Rejeb et al., 2008). Behavior abnormalities belonging to the autistic spectrum, aggression, schizophreniform psychosis have been reported in patients with PAK3 mutations (Horvath et al., 2018; Qian et al., 2020). Conversely, the role of PAK3 in schizophrenia has been investigated using pharmacological inhibitors (Hayashi-Takagi et al., 2014). Owing that cortical INs start expressing PAK3 as they enter the embryonic cortex (Cobos et al., 2007) and that neuro-psychiatric diseases including epilepsy, schizophrenia, autism, can be correlated with abnormal locations, densities and ratios of IN subpopulations (Marín, 2012), it can be hypothesized that PAK3 mutations affect brain circuit formation by disorganizing IN distribution.

PAK3 point mutations in human families all differ from each other although most of them target the catalytic domain of the protein (Qian et al., 2020). PAK3 belongs to a subgroup of the p21-activated kinase family, which can form homo- or hetero-dimers in which kinase domains are mutually inactivated (Bokoch, 2003; Combeau et al., 2012). The binding of activated (GTP linked) GTPases, RAC or CDC42, dissociates monomers and activates the kinase domain, which is thus alternatively inactive and activated in physiological conditions (Bokoch, 2003). PAKs can moreover be activated by adaptor proteins, lipids and numerous interacting proteins, such as the PIX/GIT complex that targets PAK either to focal adhesions or to the centrosome (Zhao et al., 2005).

To examine how the PAK3 kinase function influences the migration and the final distribution of INs in specific cortical areas and layers, we therefore transfected INs with PAK3 mutants locked in either a kinase activated (constitutively active PAK3-ca) or a kinase inactive (kinase dead PAK3-kd) conformation.

### PAK3 activation and PAK3 expression level

The present study aimed at characterizing the role of the PAK3 kinase function in GABAergic INs during cortical migration. Since a previous study examined the influence of *Pak3* overexpression on IN migration (Cobos et al., 2007), we also compared the expression of PAK3 mutants with the expression of a control PAK3 construct (wild-type, PAK3-wt) and with the constitutive depletion of PAK3 in *Pak3* KO INs. Whereas the morphology and dynamics of INs expressing either PAK3 mutants significantly differed from each other and from control INs expressing eGFP, neither PAK3 deleted INs nor PAK3-wt expressing INs significantly differed from control INs, suggesting that the cyclic activation/inhibition of PAK3 kinase is a critical regulator of IN migration. Noticeable differences only concerned PAK3-wt expressing INs, with a tendency for the leading process length to decrease and for the nuclear speed during fast translocations to decrease minimally. However, the cortical distribution of INs remained unchanged. Cobos and colleagues (Cobos et al., 2007) previously reported that acute *Pak3* overexpression does not affect the neuritic growth in wild type INs, whereas constitutive *Pak3* overexpression in *Dlx1-/-Dlx2-/-* INs associates with longer neurites and increased dendritic complexity. In this model, the tangential migration was moreover arrested and defaults were rescued by Pak3 siRNAs (Cobos et al., 2007). INs with either *Dlx1* or *Foxg1* deletion, that both promote *Pak3* overexpression, presented similar increase in neurite growth and dendritic complexity (Dai et al., 2014; Shen et al., 2019). Whether *Pak3* overexpression correlates with increased kinase activity has not been tested in these animal models.

In the present study, we show that INs expressing an active PAK3 mutant stopped their tangential migration as reported in *Dlx1/2 -/-* INs. However, leading processes were shortened, a default opposite to that observed in *Dlx1/2 -/-* INs. Contradictory results have also been reported in cell lines overexpressing activated PAK1: numerous protrusions and increased migration were observed in some cells (Sells et al., 1999), stable focal adhesions and impaired migration in other cell types (Kiosses et al., 1999). As suggested in cell lines, the influence of PAK activity on the neurite growth could depend on the physiological state of INs, either migrating (this study) or differentiating (Cobos et al., 2007; Dai et al., 2014; Shen et al., 2019).

Alternatively, Combeau et al. showed that PAK3 preferentially dimerizes with PAK1, which is expressed at higher levels than PAK3 in the developing brain (Combeau et al., 2012; Kreis and Barnier, 2009). Cortical GABAergic neurons likely express PAK1 in addition to PAK3 (Causeret et al., 2009). PAK3 overexpression in cortical INs could thus significantly interfere with PAK1 activation. Reciprocally, a putative “buffering” effect of PAK1 on PAK3 overexpression remains to be investigated. Analyses of *Pak3* KO and *Pak1/Pak3* double KO mice support important interactions between both isoforms (Huang et al., 2011; Meng et al., 2005).

### PAK3 controls leading process morphology and leading process orientation in INs

Our study shows that INs expressing PAK3 mutants differed from control INs and differed from each other by the morphology of their leading processes, in line with their growth cones dynamics. INs expressing PAK3-ca extended significantly shorter leading processes than control INs, and processes ended with small growth cones that frequently collapsed. By contrast, INs expressing PAK3-kd extended highly branched leading processes resulting from frequent growth cone divisions. Previous studies showed that the reorientation of cortical INs from tangential pathways to the cortical plate was accompanied by drastic morphological changes of leading processes that extended novel protrusions towards the cortical plate while they retracted tangentially oriented processes (Lysko et al., 2011; Martini et al., 2009). Despite their branched leading process, INs expressing PAK3-kd remained oriented tangentially whereas INs expressing PAK3-ca preferentially oriented radially along or between cortical plate cells (see graphical summary). These observations suggest that PAK3 activity is required to establish specific interactions between INs and cells in the cortical plate in order to promote cell turning. Signals able to maintain cortical INs in their tangential migratory streams by either attractive or repulsive activity have been identified in the developing cortex (Tiveron et al., 2006; Zimmer et al., 2011). Here we show that PAK3 activation both inhibited the tangential migration of cortical INs and promoted their radial insertion within the cortical plate, thereby controlling the tangential to radial switch of migration that allows INs to enter and colonize the cortical plate. Adhesion to cortical plate cells likely favor this switch of migration (Elias et al., 2010).

RAC and CDC42 are the two major activators of PAK kinases. Migrating cortical INs express both *Rac1* and *Rac3* that exert partly redundant functions (de Curtis, 2014). Several guidance cues able to activate Rac are moreover known to influence IN trajectories within the cortex (de Curtis, 2014; DeGeer et al., 2013; Stanco et al., 2009; Zimmer et al., 2011). Some morphological abnormalities observed in Rac1/3 deleted INs recall those of INs expressing the non activable PAK3 mutant (PAK3-kd), in particular the increased number of neurites and a very remarkable splitting of the leading process (Tivodar et al., 2015). Despite smaller growth cones, the motility of PAK3-kd expressing INs in the tangential cortical pathways remains unchanged, as previously described in *Rac* depleted INs (Tivodar et al., 2015; Vidaki et al., 2012). By contrast, PAK3-kd expressing INs hardly cross the pallium/subpallium boundary in electroporated embryonic brains, a default that recalls the impaired migration of INs from the basal to the dorsal forebrain in *Rac1/3* double KOs (Tivodar et al., 2015). Similarities between morphological and migratory defaults in INs expressing PAK3-kd and in RAC depleted INs are compatible with a role for RAC in activating PAK3 in migrating INs. Whether RAC can promote the radial re-orientation of INs by activating PAK3 is an opened question. RAC1 has been shown to regulate the morphology and dynamics of the leading process of radially migrating cells (Kawauchi et al., 2003; Konno et al., 2005). We previously showed that N-cadherin mediates interactions between migrating INs and radially oriented cells in the cortical plate (Luccardini et al., 2013). The role of PAK3 in regulating these interactions remains to be explored. Further investigations might identify N-cadherin mediated cell-cell adhesion as a molecular effector of the tangential to radial switch of migration of INs.

CDC42 has already been identified as a PAK3 activator in cortical neurons (Kreis et al., 2007). CDC42 cooperates with RAC at the leading edge of moving cells to regulate cell adhesion and actin dynamic (Etienne-Manneville, 2008; Govek et al., 2011; Lawson and Ridley, 2018). In addition, CDC42 has been shown to promote the extension and to stabilize protrusions in the developing cortex (Cappello et al., 2006). Both GTPases could cooperate or successively activate PAK3 to control and stabilize the radial orientation of the IN processes in the cortical plate.

### PAK3 activity influences centrosome positioning and nuclear movements (cell polarization/directionality)

Cdc42 is largely involved in the activation of a variety of signaling pathways regulating cell polarity, directed migration and cell adhesion (Azzarelli et al., 2014; Etienne-Manneville, 2008; Govek et al., 2011; Lawson and Ridley, 2018). In migrating cells, the polarity is controlled by the positioning of the centrosome, in font of the nucleus, at the basis of the leading process. The expression of PAK3-ca and PAK-kd in migrating INs influenced the nuclear-centrosomal distance in opposite ways. The centrosome moved forward over a long distance in the branched leading processes of INs expressing PAK3-kd, but remained close to the nucleus in INs expressing PAK3-ca. Accordingly, PAK3-kd expressing INs showed directed nuclear movements, whereas nuclear movements in INs overexpressing PAK3-ca showed frequent direction reversals, identifying the nuclear centrosomal distance as the master regulator of cell polarity and of sustained directionality in these neurons. The precise role of PAK3 in controlling the centrosome position in migrating INs remains unclear. The PIX/GIT complex can recruit and activate PAK at the centrosome (Zhao et al., 2005). Cdc42 has been shown to accumulate in the centrosomal region of migrating neurons (Konno et al., 2005) but its capacity to activate PAKs near the centrosome is not known. It has been shown in the cerebellum that cerebellar granule cells invalidated for Cdc42 or overexpressing a dominant negative Cdc42 show polarity defaults and impaired migration junctions with Bergman glia associated with phosphorylation changes in PAK proteins. However, the expression level of the protein complex that regulates centrosomal movements in these neurons remained unchanged (Govek et al., 2018; Solecki et al., 2004). POSH, an activator of Rac1 has also been localized in the rostral swelling of radially migrating cortical neurons where the centrosome positions before nuclear translocation (Yang et al., 2012). Alternatively, PAK proteins could control the dynamics of microtubules by blocking the microtubule depolymerizing activity of stathmin through phosphorylation (Kwon et al., 2020), thereby regulating the length and stability of microtubule bundles linking the centrosomal swelling to the nuclear compartment and to the leading process (Baudoin et al., 2012; Solecki et al., 2009).

By monitoring a non-muscle myosin IIB-GFP (Myosin 2B-GFP) fusion protein (Bao et al., 2007) in PAK3-ca and PAK-kd expression INs, we showed that PAK3 kinase activity significantly altered the subcellular distribution of Myosin 2B in migrating cortical INs and regulated the acto-myosin dependent contractility of the cell body (Bellion et al, 2005; Godin et al., 2012). Myosin 2B is a non-muscle myosin II isoform that defines the non-protrusive parts of migrating cells -nuclear compartment and trailing edge- and regulates nuclear dynamics (Vicente-Manzanares et al., 2009). Myosin 2B seems functionally predominant in tangentially migrating neurons (Ma et al., 2004). The regulatory light chain (RLC) of the non-muscle myosin II activates both Myosin 2B and Myosin 2A but different regulatory mechanisms seem involved (Sandquist et al., 2006). PAK kinases regulate myosin contractility by activating the RLC on one hand, and the myosin light chain kinase (MLCK) that represses RLC on the other hand (Sanders et al., 1999). Despite sustained efforts (JV Barnier & V. Rousseau, data not shown), we could not show any effect of PAK3 on the phosphorylation state of neither RLC in Myosin 2B, nor MLCK. Whether PAK3 activity indirectly interferes with the RhoA/ROCK signaling pathway known to control the contractility of the acto-myosin cytoskeleton in migrating INs (Godin et al., 2012) remains to be explored.

Whereas acto-myosin filaments accumulated at the cell rear and in the centrosomal region of PAK3-kd expressing INs with long nuclear-centrosomal distance, PAK3-ca expressing INs, on the contrary, comprised fainter myosin filaments associated with shorter nuclear-centrosomal distances. This suggests that PAK3 activity either displaced and/or disassembled Myosin 2B filaments by unknown mechanisms and that Myosin 2B contractility is required to maintain long nuclear-centrosomal distance in migrating INs. Myosin 2B moreover concentrated in growth cones of PAK3-kd expressing INs before splitting, suggesting that Myosin 2B activity in the leading process contributed to growth cone splitting.

## CONCLUSION

In the present study, we have identified PAK3 as a kinase whose activity in cortical INs interrupts the tangential migration and promotes the radial re-orientation and migration within the cortical plate. All together, these results suggest that cortical plate colonization by INs relies on the capacity for PAK3 kinase to switch from an inactive to an active state. PAK3 activity disrupts the polarized morphology required for the tangential migration of INs by i) preventing the formation and stabilization of branches in which the centrosome migrates forward, and ii) disrupting acto-myosin filaments and preventing them to maintain the centrosome a long distance ahead the nucleus. Multiple signals could trigger PAK3 activation in the tangential cortical pathways. PAK3 mutations targeting the kinase domain of the protein, most frequently found in patients with ID and cortical malformations, should therefore severely impair the migration of cortical interneurons in infants and the formation of normal cortical circuits. PAK3 depletion seems much less deleterious for the cortical targeting of INs than PAK3 kinase alteration, likely due to compensatory effects by other regulatory complexes.

## Acknowledgements

This work was supported by Institut National de la Santé et de la Recherche Médicale (INSERM), Centre National de la Recherche Scientifique (CNRS), Sorbonne University, Paris-Saclay University. Agence Nationale de la Recherche (ANR, grant MRGENES to C.M. and J.-V.B., and grant MIGRACIL to C.M.), Fondation pour la Recherche sur le Cerveau (grant R11080DD to C.M.), Fondation J.Lejeune (grant R14108DD to C.M.).and Japan Society for the Promotion of Science (JPSP). L.V. was supported by a thesis fellowship from Ministère de la Recherche et Technologie, a 6 month fellowship from Fondation pour la Recherche Médicale, and an INSERM-JPSP travel grant. P.-S.L. was supported by ANR (grant MRGENES).

Professor Miyzaki and Dr Gauthier-Rouvière are acknowledged for the gift of expression vectors. We thank Véronique Devignot and Matsutoshi Yanagida for their assistance with the project, A. Hanauer, C. Vaillend and all members of the Métin’s Team for constructive discussions. We thank Stéphane Robin for statistical analyses and Melody Atkins for english revision. We gratefully acknowledge the Imaging plateform facility of the Fer à Moulin Institute for the use of their microscopes, and the animal facility of the Fer à Moulin Institute for animal care and breeding.

## LEGENDS

**Movie 3A1 (related to Fig. 3-A1) : Migration of MGE cells expressing eGFP in an organotypic cortical slice.** The electroporated MGE explant has been grafted in the lateral cortex. It releases eGFP(+) cells that migrate toward the medial cortex. Time-lapse between frames is 10 minutes (sequence of 10 hours, 15 frames per second).

**Movie3A2 (related to Fig. 3-A2): Migration of MGE cells expressing eGFP-PAK3-ca in an organotypic cortical slice.** The electroporated MGE explant has been grafted in the lateral cortex. It releases eGFP(+) cells that migrate in the cortical plate and below the cortical plate. Time-lapse between frames is 10 minutes (sequence of 10 hours, 15 frames per second).

**Movie3A3 (related to Fig. 3-A3) : Migration of MGE cells expressing eGFP-PAK3-kd in an organotypic cortical slice.** The electroporated MGE explant has been grafted in the lateral cortex. It releases eGFP(+) cells that migrate toward the medial cortex. Time-lapse between frames is 10 minutes (sequence of 10 hours, 15 frames per second).

**Movie4B (related to Fig. 4-B): Migration of MGE cells expressing eGFP in a co-culture.** The electroporated MGE explant has been cultured on a substrate of dissociated cortical cells. It is positioned on the left side of the movie. Time-lapse between frames is 3 minutes (sequence of 4.5 hours, 15 frames per second).

**Movie4C (related to Fig. 4-C): Migration of MGE cells expressing eGFP-PAK3-ca in a co-culture.** The electroporated MGE explant has been cultured on a substrate of dissociated cortical cells. It is positioned on the left-bottom corner of the movie. Time-lapse between frames is 3 minutes (sequence of 5.5 hours, 15 frames per second).

**Movie4D (related to Fig. 4-D): Migration of MGE cells expressing eGFP-PAK3-kd in a co-culture.** The electroporated MGE explant has been cultured on a substrate of dissociated cortical cells. It is positioned on the left side of the movie. Time-lapse between frames is 3 minutes (sequence of 3.8 hours, 15 frames per second).

**Movie5E (related to Fig. 5-E): Dendrogram reproduces the dynamic transformations of growth cones at the end of neurite branches over time.** Arrowheads are migrating growth cones, bars represent growth cone collapse, and yellow branches are growth cone splitting. Time-lapse between frames is 3 minutes (sequence of 1.0 hour, 10 frames per second).

**Movie6A2 (related to Fig. 6-A2): Migration of a MGE cell co-expressing eGFP and a red centrosomal marker (pericentrin-mKO1) in a co-culture.** Time-lapse between frames is 3 minutes (sequence of 2.25 hours, 15 frames per second).

**Movie6B1 (related to Fig. 6-B1): Migration of a MGE cell co-expressing eGFP-PAK3-kd and a red centrosomal marker (pericentrin-mKO1) in a co-culture.** Time-lapse between frames is 3 minutes (sequence of 5 hours, 15 frames per second).

**Movie6B2 (related to Fig. 6-B2): Migration of a MGE cell co-expressing eGFP-PAK3-kd and a red centrosomal marker (pericentrin-mKO1) in a co-culture.** Time-lapse between frames is 3 minutes (sequence of 3 hours, 15 frames per second).

**Movie7A (related to Fig. 7-A): Migration in a co-culture of a Myosin 2B-GFP expressing MGE cell electroporated with mRFP**. A MGE explant from a Myosin 2B-GFP embryo has been electroporated and cultured on wild-type dissociated cortical cells. Time-lapse between frames is 1 minute (sequence of 1.5 hour, 10 frames per second).

**Movie7C1 (related to Fig. 7-C1): Migration in a co-culture of a Myosin 2B-GFP expressing MGE cell electroporated with mRFP-PAK3-ca**. A MGE explant from a Myosin 2B-GFP embryo has been electroporated and cultured on wild-type dissociated cortical cells. Time-lapse between frames is 1 minute (sequence of 1.85 hour, 10 frames per second).

**Movie7C2 (related to Fig. 7-C2): Migration in a co-culture of a Myosin 2B-GFP expressing MGE cell electroporated with mRFP-PAK3-kd**. A MGE explant from a Myosin 2B-GFP embryo has been electroporated and cultured on wild-type dissociated cortical cells. Time-lapse between frames is 1 minute (sequence of 1.8 hour, 10 frames per second).

---

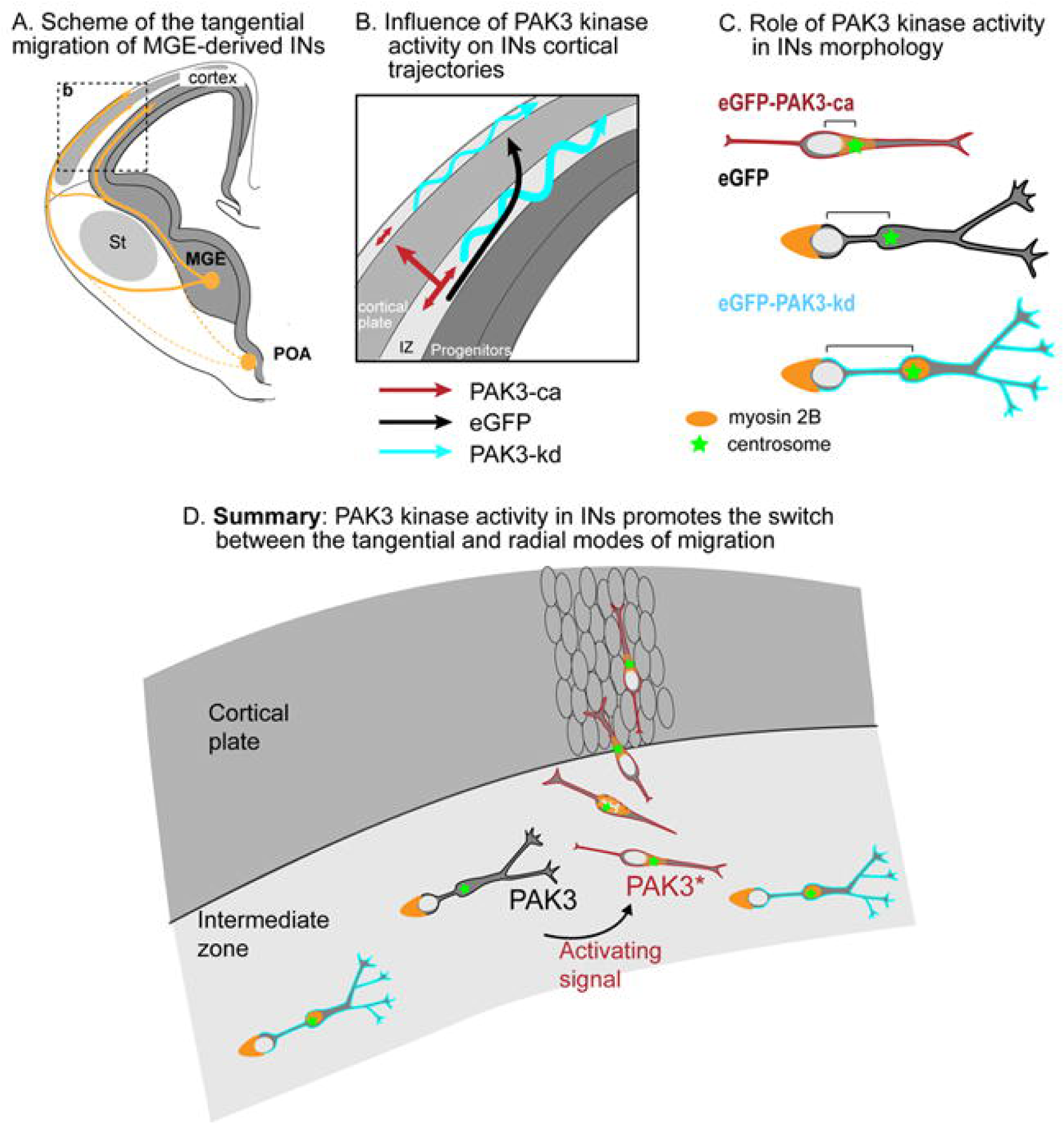

