## Supplemental Figures (1-7) Table1 Methods for "PAK3 controls the tangential to radial migration switch of cortical interneurons by coordinating changes in cell shape and polarity"

**Supplemental information**

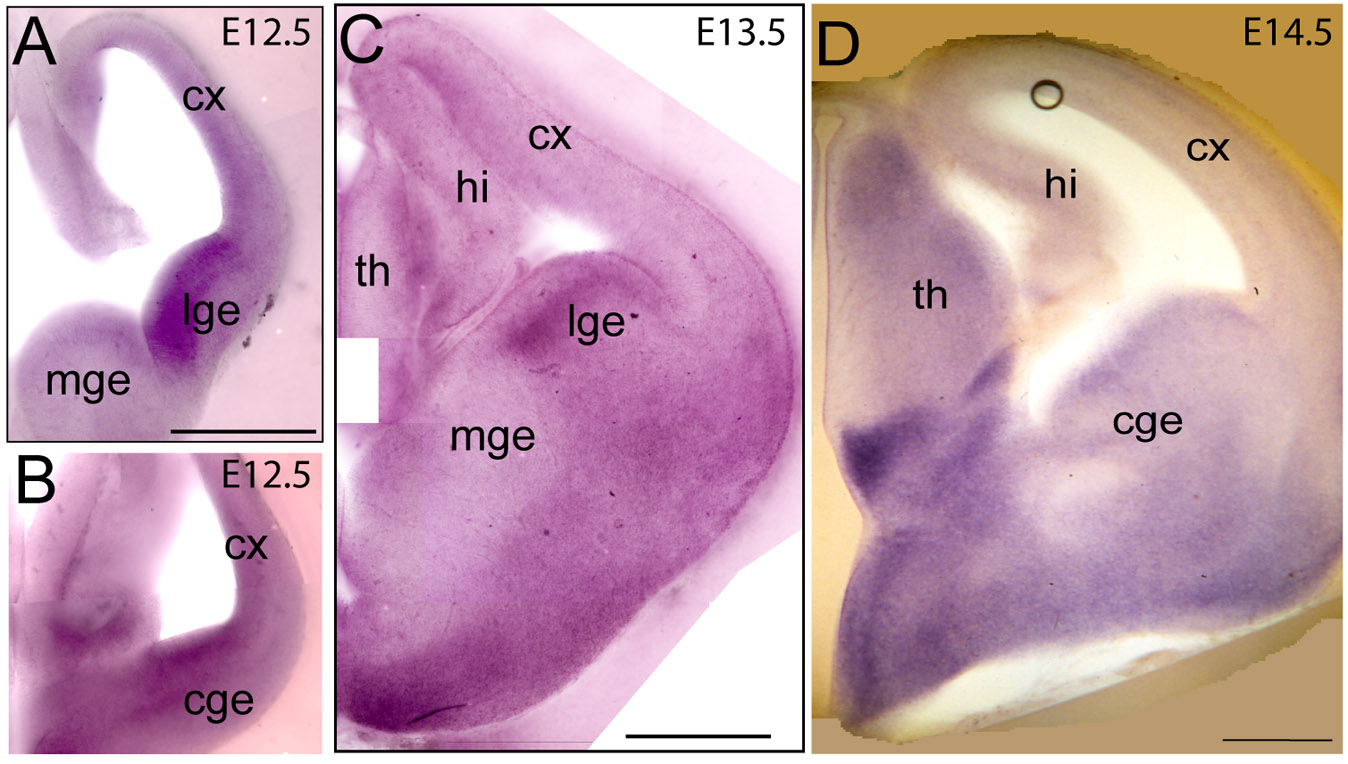
**Figure S1 (related to Fig.1): Pak3 expression in the mouse telencephalon at embryonic stages 12.5, 13.5 and 14.5.**

Frontal 150 μm-thick sections of embryonic mice forebrains at E12.5 (**A**, medial section, **B** caudal section), E13.5 (**C**, medial section), and E14.5 (**D**, caudal section) were hybridized with an anti-Pak3 probe. A strong signal is observed in the proliferative zone of the lateral (lge) and caudal (cge) ganglionic eminences, but not in the medial ganglionic eminence (mge). Scale bars: (A,C,D) 500 μm.

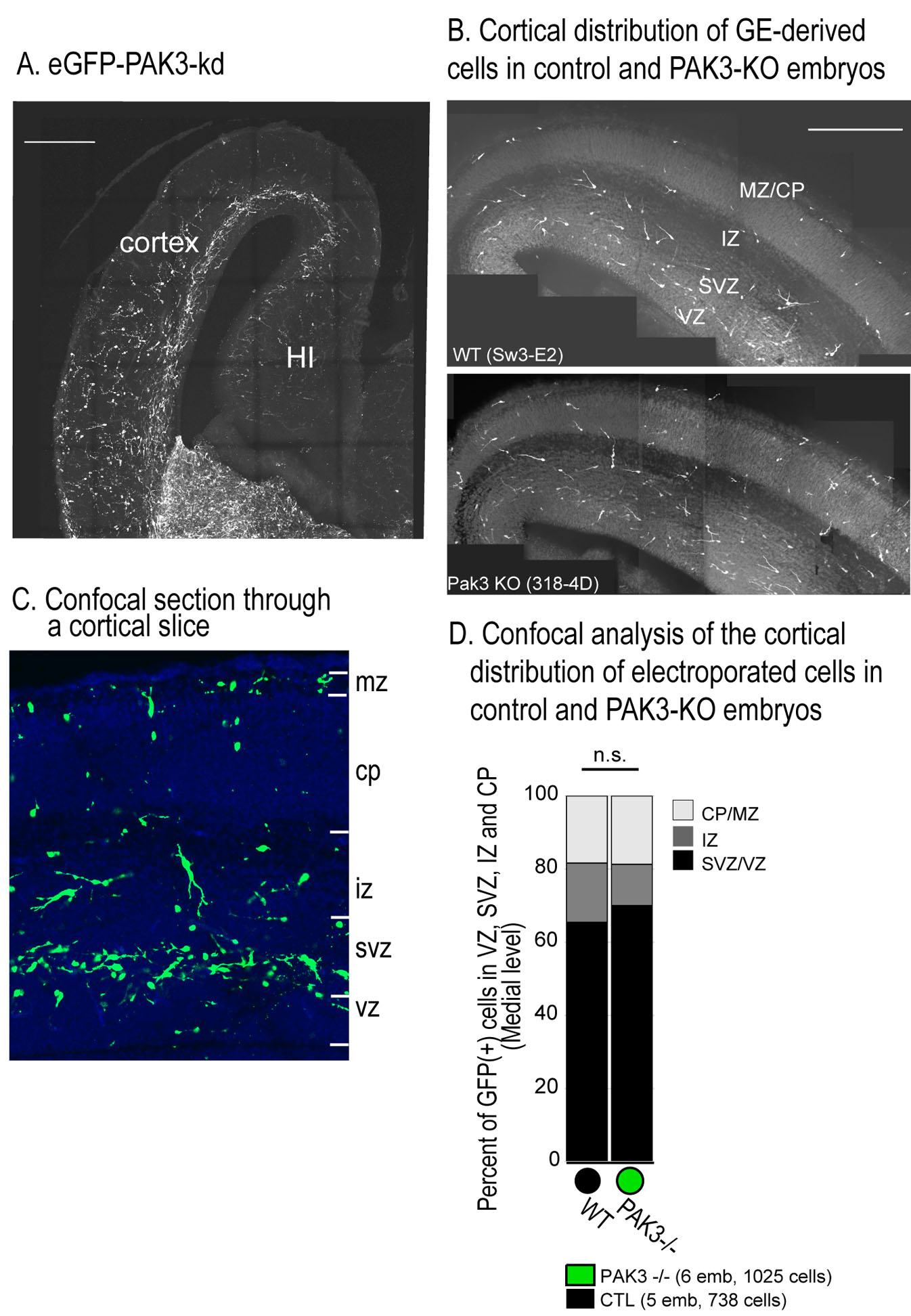

**Figure S2 (related to Fig.2): Cortical distribution of interneurons expressing the PAK3-kd mutant or knocked out for *Pak3*.**

**A.** Picture of a 50μm-thick coronal section through the cortex of an E16.5 embryo electroporated at E12.5 with the eGFP-PAK3-kd construct. A large stream of GE-derived electroporated cells is present along the ventricular surface. HI, hippocampus. Scale bar: 300 µm. **B.** Photomicrographs illustrate the cortical distribution of GE-derived cells electroporated at E12.5 with eGFP in E16.5 wild type (WT) (top) and *Pak3*-KO (bottom) embryos. Coronal sections are 50 μm thick. Scale bar: 300 μm **C**. The cortical distribution of eGFP+ cells was analyzed in DAPI counterstained sections using a confocal microscope. Quantification was performed every 2 µm in a 30 µm-thick section. **D**. Histogram comparing the distribution within the layers of a dorsal cortical sector (positioned as the dotted line rectangle in Fig.2C1) of cells electroporated with eGFP in the GE of wild type (swiss) and *Pak3*-KO embryos. Distributions do not significantly differ in a Chi2 test. cp, cortical plate; iz, intermediate zone; mz, marginal zone; svz, subventricular zone; vz, ventricular zone.

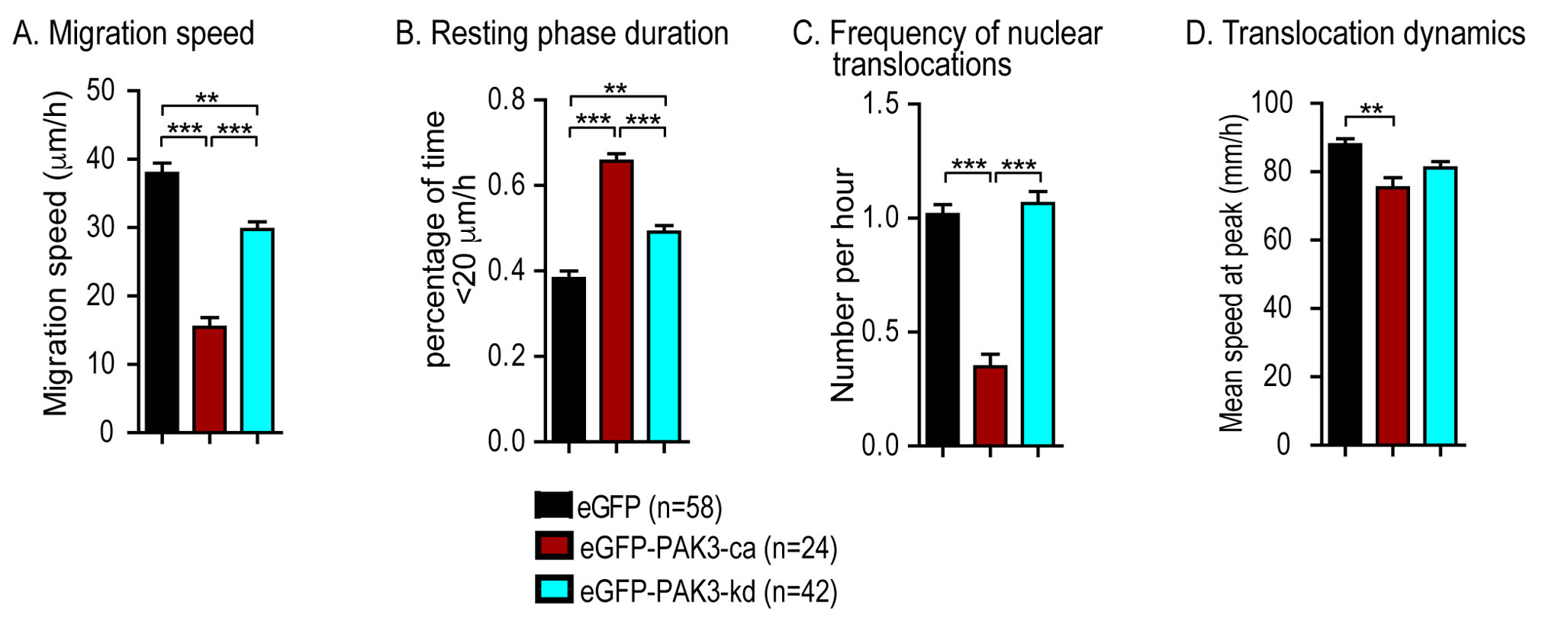

**Figure S3 (related to Figure 3): Cell body dynamics of MGE cells grafted in cortical slices.**

**A.** Histogram showing the mean migration speed of eGFP- (black bars), eGFP-PAK3-ca- (red bars) and eGFP-PAK3-kd- (blue bars) expressing MGE cells that migrated in the deep stream (red trajectories in Fig.3B1,B2, B3). **B, C, D.** Histograms analyze the saltatory progression of MGE cells in grafted slices. **B** shows the duration of resting phases as a percentage of the recording duration. **C, D**. Slices were imaged every 10 minutes. The threshold speed to define nuclear translocations was 50 μm/hour. Histograms give the frequency of nuclear translocations (C) and the mean cell body speed at the peak (D). Statistical significance was assessed by Kruskal-Wallis tests (p<0.0001) and Dunn’s Multiple comparison tests (p<0.001 and p<0.01 as indicated on histograms).

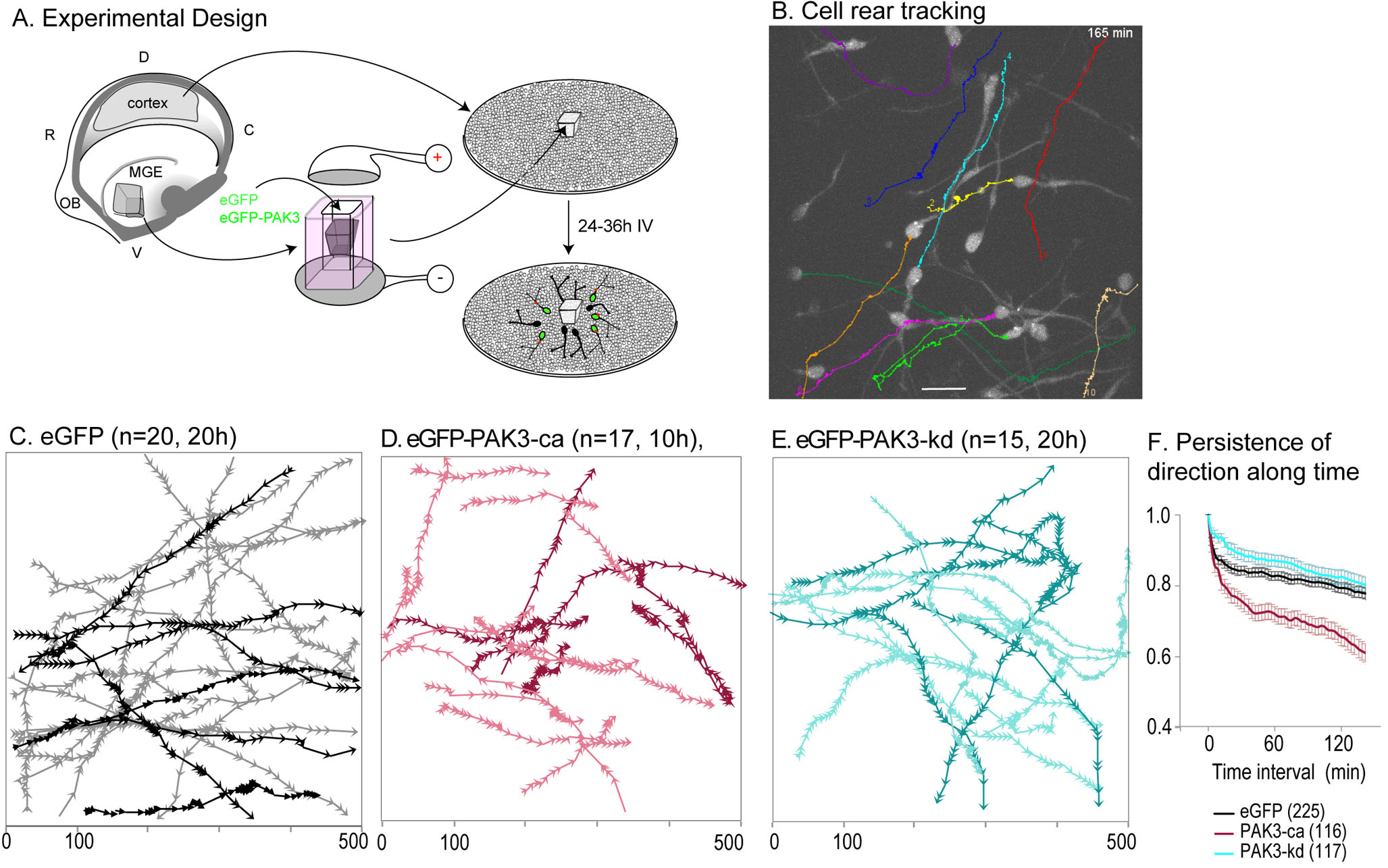

**Figure S4 (related to Figure 4): Analysis of MGE cells trajectories in co-cultures.**

**A.** Scheme of the experimental design. MGE explants (green) were dissected out of telencephalic vesicles at E13.5, electroporated and placed on a substrate of E13.5 wild type dissociated cortical cells. **B.** eGFP MGE cells migrating away from explants on the substrate of cortical cells were imaged by time-lapse video-microscopy. The rear of cells was tracked manually using the MTrakJ plugin of ImageJ on movies to reconstitute cell trajectories (colored traces). **C, D, E.** Frames are representative examples of trajectories of cells electroporated with eGFP, eGFP-PAK3-ca and eGFP-PAK3-kd and recorded in a unique field of view at the migration front. Arrowheads indicate the position of the cell body at each time point, and the direction of movement of the cell. **F.** Curves show the temporal evolution of the direction persistence index

(Gorelik and Gautreau, 2014) of MGE cells transfected with eGFP (black), eGFP-PAK3-ca (red) and eGF-PAK3-kd (blue). eGFP-PAK3-ca expressing MGE cells (red curve) are significantly less directional than control MGE cells, whereas eGFP-PAK3-kd expressing MGE cells (blue curve) are significantly more directional than control MGE cells until 80 minutes.

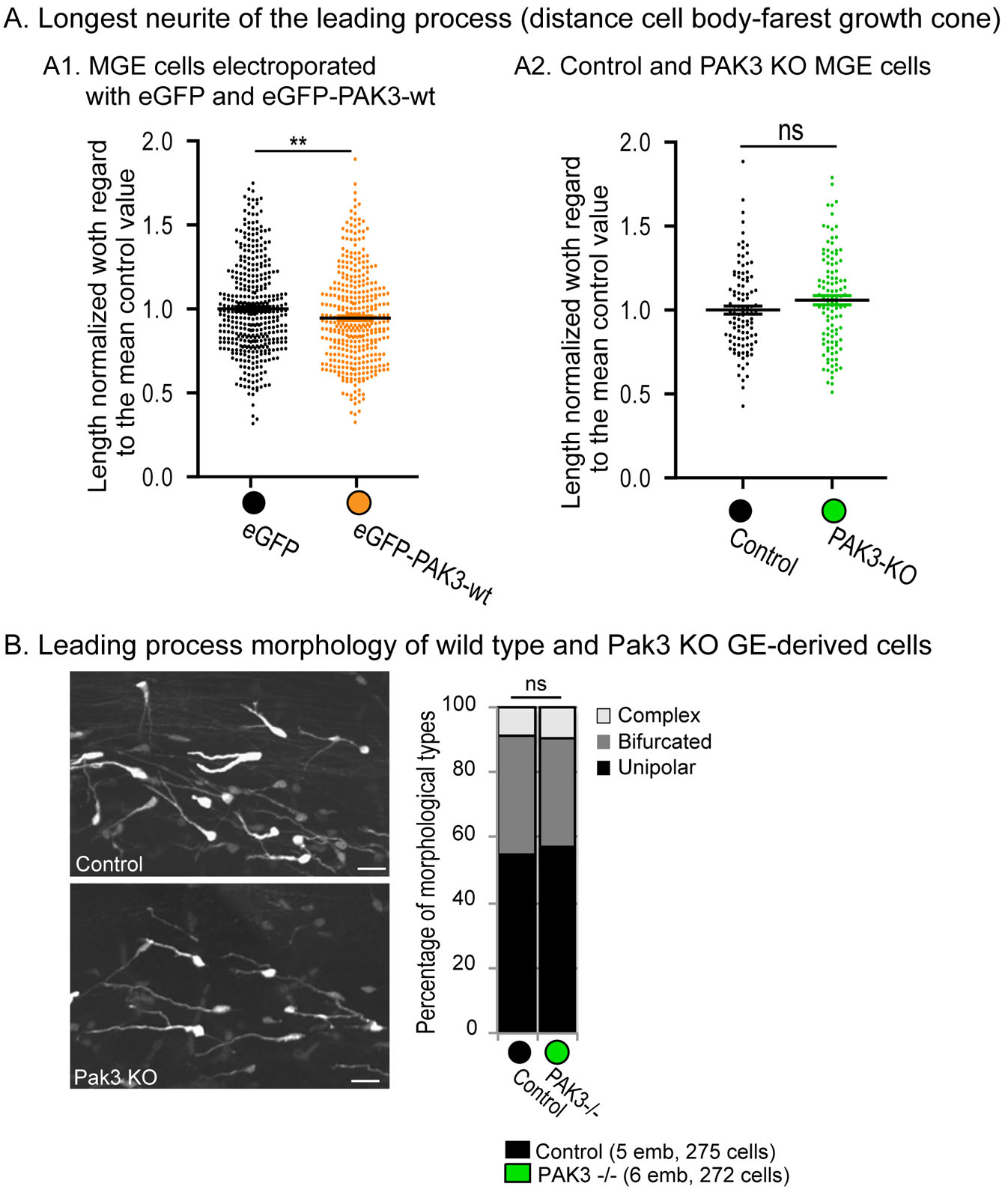

**Figure S5 (related to Figure 5): Analysis of the leading process morphology in MGE cells expressing the PAK-wt construct and in *Pak3*-KO MGE cells.**

**A.** Measure of the longest neurite in leading processes of MGE cells migrating away from their explant of origin on dissociated cortical cells. A1. Comparison between MGE cells electroporated with eGFP and eGFP-PAK3-wt plasmids. A2. Comparison between control (WT) and *Pak3*-KO MGE cells electroporated with eGFP. Statistical differences were assessed by Mann Whitney T-tests (**, p=0.006; n.s., p=0.135) **B.** eGFP was electroporated at E12.5 in the GE of control and PAK3-KO embryos. Electroporated embryos were fixed at E16.5 and eGFP(+) GE-derived cells imaged in the cortex on coronal sections by confocal microscopy. The branching pattern of eGFP(+) WT and eGFP(+) *Pak3*-KO GE-derived cells did not differ by a Chi2 test.

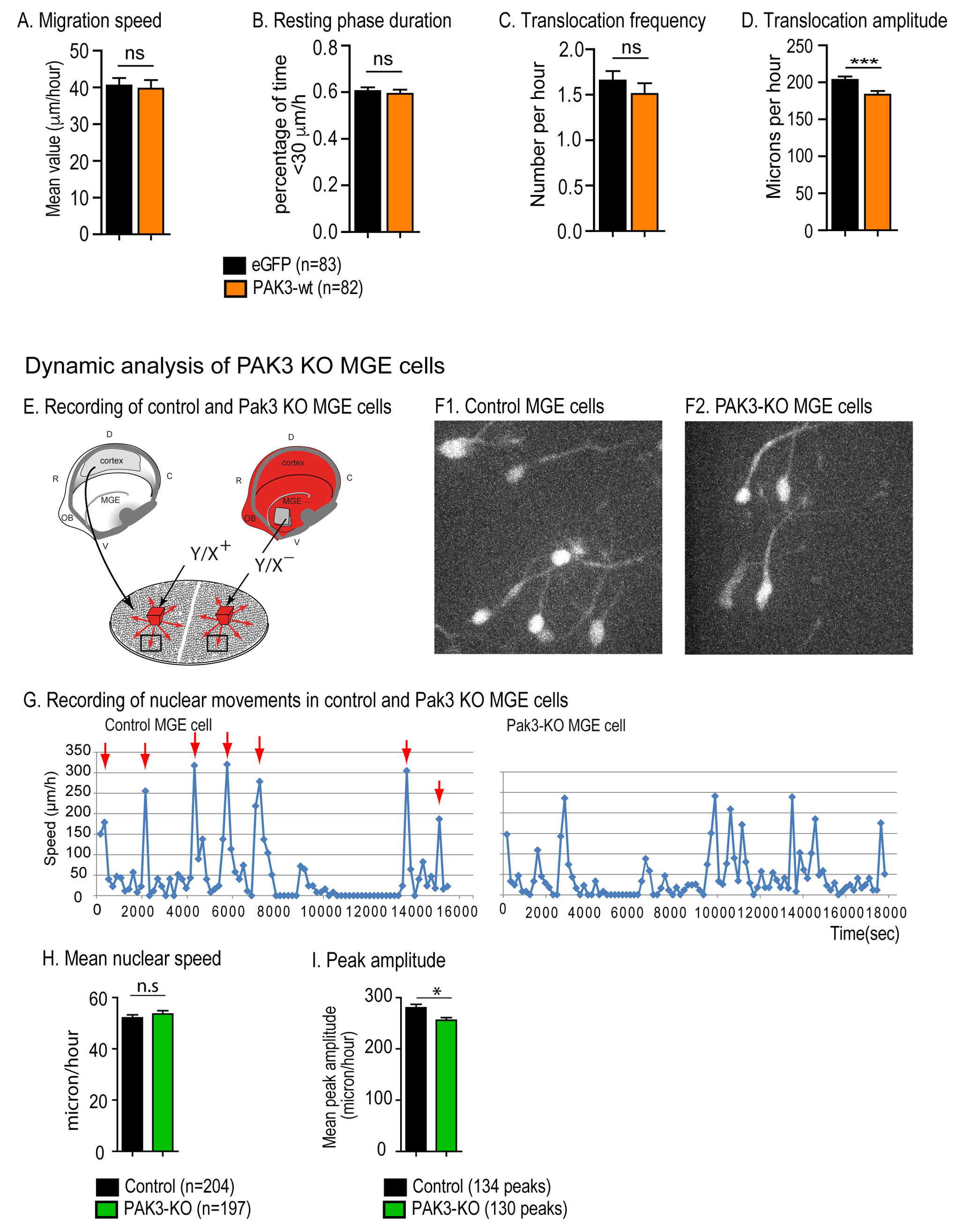

**Figure S6 (related to Figure 6): Analysis of nuclear movements in PAK3-wt expressing MGE cells and in *Pak3* KO MGE cells in co-cultures.**

**A, B, C, D.** Mean migration speed and nuclear dynamics in eGFP (black bars) and eGFP-PAK3-wt (orange bars) expressing MGE cells that migrated on dissociated cortical cells in co-cultures (see Fig.S4-A). Statistical significance was assessed by Mann-Withney tests (*** p<0.001). **E, F1, F2, G, H, I.** Nuclear movements in control and PAK3 KO MGE cells migrating on wild type dissociated cortical cells. Migrating cells were analyzed by time-lapse video-microscopy in co-cultures as illustrated in scheme **E**. **F1** and **F2** are two movie frames. Scale bar, 20 μm. Graphs in **G** compare the nuclear movements in two representative control and PAK3-KO MGE cells. **H, I.** Histograms compare the mean nuclear speed (H) and the mean amplitude of nuclear translocations (> 100 μm/hour) (I) in control (black bars) and PAK3-KO (green bars) MGE cells. Statistical significance was assessed by Mann-Withney tests (*, p=0,0110).

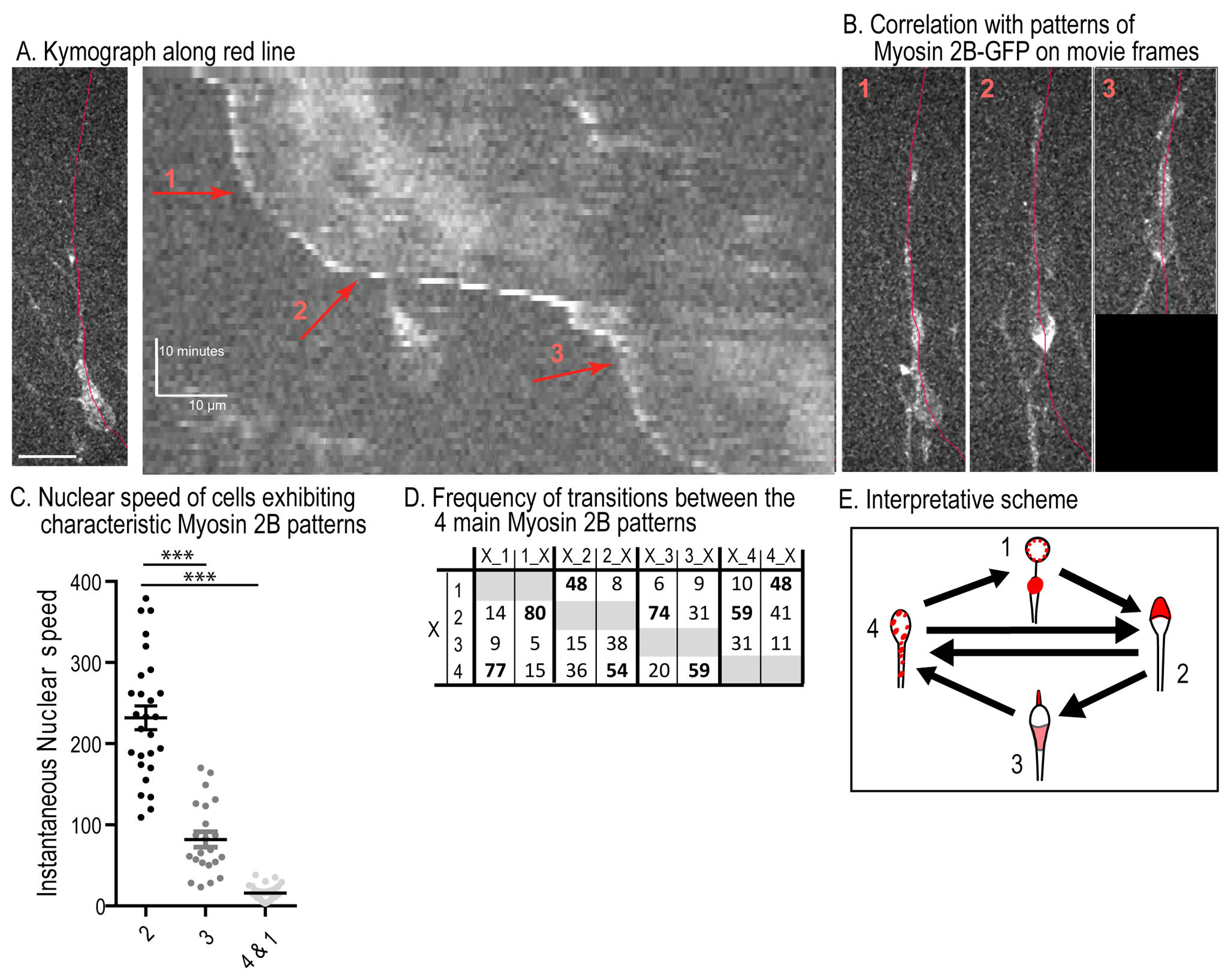

**Figure S7 (related to Figure 7): Patterns of Myosin 2B distribution during the migration cycle in MGE cells.**

**A.** Kymograph shows the temporal evolution of Myosin 2B-GFP distribution along the red line in the migrating MGE cell shown on the left. Scale bars on kymographs show time (y=10 minutes) and distance (x= 10 microns). **B.** Myosin 2B-GFP distributions corresponding to the patterns 1,2,3 defined in Fig.7 are identified on the kymograph. **C.** Histogram shows the speed of cell rear calculated from kymographs and expressed as a function of the Myosin 2B-GFP patterns defined in Fig.7. **D.** Transitions between the 4 patterns of Myosin 2B-GFP were identified on movies and their frequencies calculated. Statistical significance was assessed using One-way ANOVA (p<0.0001) and post-hoc Dunn’s tests (results indicated on graph). **E.** Arrows on the scheme figure the most frequent transitions between Myosin 2B-GFP patterns in control MGE cells during migration.

**Table S1 (related to Table 1): Number of wild type and *Pak3* KO embryos electroporated with an eGFP plasmid in the ganglionic eminences (GEs) at E12.5.** Proportion of embryos that survived until E16.5 and in which electroporated GE cells reached the cortex.

| Plasmid |  | Number of electroporated embryos | Nber of surviving embryos | Nber of surviving embryos with cortical streams of electroporated GE cells | Mean number of cells per embryo (lateral + dorsal sectors) |
| --- | --- | --- | --- | --- | --- |
| pCAG-eGFP | Swiss | 20 | **15** (75%) | 7 (47% of surviving) | 148 +41 |
| pCAG-eGFP | *Pak3* KO | 18 | **14** (78%) | 9 (64% of surviving) | 171 + 70 |

METHODS

**Mice**

Mice were housed and mated in a conventional animal facility according to European guidelines. Experiments performed in the present study have been validated and approved by the Ethical committee Charles Darwin (C2EA-05, authorized project 02241.02). Midday of the day of vaginal plug formation was considered E0.5.

Mice from the following strains were used at the adult stage: *Swiss* (Janvier, France) and *Pak3* KO mutant mice (Meng et al., 2005).

Mice embryos were produced by crossing wild type or genetically modified adults: *Swiss* mice (Janvier, France or Charles River breeders), transgenic (CAG-mRFP1) mice that ubiquitously express the RFP (generous gift from N. Hadjantonakis, (Long et al., 2005), *Pak3* KO mice, and mutant mice expressing the GFP-tagged full length NMHC II-B (Myosin 2B-GFP (Bao et al., 2007).

**Plasmids**

pCAG-eGFP was a generous gift from Prof Miyzaki (Osaka University, Japan) and pCAG-mRFP from Prof F. Murakami (Osaka University, Japan). They were used at a final concentration 0.5 µg/µL for *in vitro* and *in utero* electroporation.

*Construction of eGFP-PAK3 and mRFP-PAK3 plasmids.*

A polylinker (P1, P2) was inserted in the EcoR1 site of the pCAG vector (gift of Prof Miyzaki, Osaka University) to generate a pPaGi vector with unique sites BspEI, XhoI, ClaI, EcoRI. GFP-PAK3 fragments were amplified with P3, P4 primers, to add BspeI and ClaI sites to the following plasmids: pGFP-PAK3a-WT,-K297L,-T421E (Kreis et al., 2007). Fragments were then sub-cloned in the pPaGi vector by double digestion BspeI/ClaI, to obtain the following plasmids: pPaGi-GFP-PAK3a-WT to express wild type PAK3-GFP fusion protein (eGFP-PAK3-wt); pPaGi-GFP-PAK3a-K297L to express kinase-dead mutant PAK3-GFP (eGFP-PAK3-kd); pPaGi-mRFP-PAK3a-T421E to express constitutively active mutant PAK3-GFP (eGFP-PAK3-ca).

The mRFP fragment was amplified from a mRFP-actin plasmid (Gift from Cécile Gauthier-Rouvière) with primers (P5, P6) that add HindIII and BamHI sites. It was then introduced in the pcDNA3 vector digested with HindIII/BamHI to result in the pcDNA3-mRFP plasmid. BamHI-XbaI fragments of PAK3 constructs were obtained from plasmids pcDNA3-HA-PAK3a-WT, K297L, T421E (Kreis et al., 2007; Rousseau, 2002) and introduced in the pcDNA3-mRFP vector by BamHI-XbaI digestion, to obtain pcDNA3-mRFP-PAK3a-WT, K297L, and T421E.

The mRFP-PAK3 fragments were amplified with primers P7, P4 to add BspeI and ClaI sites to plasmids pcDNA3-mRFP-PAK3a-WT, K297L, and T421E. The fragments were sub-cloned in the pPaGi vector by double digestion BspeI-ClaI, to generate the following plasmids: pPaGi-mRFP-PAK3a-WT (RFP-PAK3-wt); pPaGi-mRFP-PAK3a-K297L (RFP-PAK3-kd); pPaGi-mRFP-PAK3a-T421E (RFP-PAK3-ca).

**Primers:**

P1 : 5’-AATT CTCCGGACTCGAGAATCGAT- 3’

P2 : 5’ -AATTA TCGATTCTCGAGTCCGGAG- 3’

P3 : 5’ –CGGCTCCGGAGCCACCATGGTCAGCAAGGGCGAGGA- 3’

P4 : 5’ –CGCATCGATTCTAACGGCTACTGTTC- 3’

P5 : 5’ –AAGCTTGCCACCATGGCCTCCTCCGAG- 3’

P6 : 5’ –GGATCCGTGGCGCCGGTGGAGTGGCGGCC- 3’

P7 : 5’ -CGGCTCCGGAGCCACCATGGCCTCCTCCGAGGACGT- 3’

Before electroporation, constructs were diluted in PBS with 0.01% Fast Green at a final concentration of 1µg/µL. For *in utero* electroporation, constructs were diluted at 1.5µg/µL.

**Cultures and *in vitro* electroporation.**

Brains and MGE explants of wild-type or transgenic/mutant embryos were collected in cold PBS at embryonic day E13.5 and dissected in cold Leibovitz medium (Invitrogen). Entire MGE explants were placed in a small well of 3% agar. A plasmid diluted in PBS and 0.01% Fast Green at a final concentration indicated above was microinjected within the explant. The explants were then placed on a petri dish equipped with electrodes (Nepagene, Sonidel, UE) and electroporated with one pulse (100 V, 5 msec) delivered by a BTX electroporator (see also (Baudoin et al., 2012) for more procedure details). Electroporated explants were then cultured for 5-6 hours in F12/DMEM medium with 10% calf serum in an incubator (37°C, 5% CO2) for “recovery”.

Electroporated MGE explants were divided into smaller pieces and placed either on dissociated cortical cells or grafted into organotypic slices. Slices were prepared from E14.5 embryonic brains embedded in 3% type VII agar and sectioned coronally using a manual slicer into 150 or 200µm thick sections. Slices were thereafter transferred in Millicell chambers (Merck Millipore) for culture.

Co-cultures used for live imaging were prepared in petri dishes equipped with a glass coverslip. Slices were transferred in a petri dish equipped with a glass coverslip before imaging. The culture medium was replaced by a culture medium of same composition but without phenol red one hour before imaging.

**Videomicroscopy**

*Co-cultures*

Time-lapse imaging was performed with an inverted microscope equipped with a spinning disk (Leica DMI4000) and with a temperature-controlled chamber. Cells were observed using either a x40 or a x63 immersion objective. Stacks were captured with a Coolsnap HQ camera (Roper Scientific, USA) every 3 or 5 min for up to 10 hr. Control and mutant MGE cells were recorded at the same time under the same acquisition conditions.

*Organotypic slices cultures*

Time-lapse imaging of grafted organotypic slices cultured for 24hours was performed with an epifluorescence macroscope (MVX10 Olympus) equipped with a temperature-controlled chamber at 34-35°C using a x2x3.2 (or a x2x4) objective. Pictures were acquired with a time interval of 5 or 10 minutes for 21 to 24hours.

Acquisitions were controlled using Metamorph software (Roper Scientific, USA).

***In utero* electroporation.**

Ganglionic eminence directed *in utero* electroporation was performed as described in (Luccardini et al., 2013) on E12.5 swiss embryos or on E13.5 PAK3 KOs embryo. Glass capillaries (Narishige, Japan) were pulled and calibrated for 2-3 μl plasmid injections. After DNA injection into the lateral ventricle, 5 pulses separated by 950 ms were applied at 60V for 50 ms at an angle of 30–60° from the horizontal plane (electrodes BEX LF650P2; BTX electroporator). Embryos were allowed to develop *in utero* for 3 to 4 days and sacrificed at E16.5. Brains were collected in cold PBS and a piece of each embryo was collected for immediate PAK3 genotyping. Brains were immediately fixed in cold 4% (wt/vol) paraformaldehyde (PFA) in 0.12 M phosphate buffer (pH 7.4) overnight. Brains were sectioned in the coronal plane at a thickness of 50 μm.

In each electroporated embryo, the cortical section containing the largest number of transfected cells was selected and transfected cells counted within dorsal and lateral sectors of same width. In both sectors, we moreover analyzed the distribution of cells in the embryonic cortical layers.

**Immunohisto- and cyto-chemistry**

Embryonic brains were dissected in cold Leibovitz medium (Invitrogen) and fixed by immersion in cold 4% (wt/vol) PFA in 0.12 M phosphate buffer (pH 7.4) overnight. Then, brains were either embedded in 3% agarose and coronally sectioned with a vibratome at 50µm, or cryoprotected in 10% sucrose in 0.12 M phosphate buffer (pH 7.4) and coronally sectioned at 20 μm with a cryostat.

Adult brains perfused with 4% PFA and cryoprotected were coronally sectioned at 40 μm with a cryomicrotome. Organotypic slices were fixed by immersion in cold 4% PFA.

Co-cultures were fixed at least 3 hours in 4% PFA/ 0.33 M sucrose in 0.12M Phosphate Buffer at 4°C for GFP staining. 0.1% picric acid was added to this fixation mix for PAK3 staining.

Cryostat and floating brain sections as well as fixed cultures were preincubated for two hours respectively in PGT (PBS; gelatin 2g/L; 0,25% (wt/vol) Triton X-100) and PBT (PBS; 0.25% Triton X-100) with 10% normal goat serum. Organotypic brain slices grafted with electroporated explants were pre-incubated in PGT for more than 5 hours before GFP amplification. Tissues or cells were then incubated overnight at 4°C in primary antibodies respectively diluted in PGT or PBT with 2% normal goat serum and 1% bovine serum albumin (BSA, Sigma-Aldrich). After 3 rinses with PBT, sections were incubated 1h30 with secondary antibodies diluted in PBT at 1/400. Brain sections were extensively washed in PBS after antibody incubation.

The following primary antibodies were used: chicken anti GFP (1:5000 Aves Lab), rabbit anti GFP serum (1:1000 Invitrogen), a rabbit polyclonal anti-PAK3 serum (rb-211-PAK3, Combeau et al, 2012, 1/100) was used for immunohistofluorescence experiments and a mouse monoclonal anti-PAK3 (clone 3A12, 1/100, Sigma-Aldrich) was used for immunohistochemistry experiments on sections pretreated with 1%H2O2 in PBS. Primary antibodies were revealed by immunofluorescence with the appropriate Alexa dye (Molecular Probes) or Cy3 (Jackson laboratories) conjugated secondary antibodies diluted in PBT (1:400). F-actin was detected with phalloidin coupled to alexa Fluor (1/40 in PBT, 20 minutes, Invitrogen). Bisbenzimide (1/5000 in PBT, Sigma) was used for nuclear counterstaining. Immunohistochemistry was performed using biotinylated goat anti-mouse IgG (Vector Laboratories, Burlingame, CA). The biotinylated antibodies were detected using the Vectastain ABC Elite Kit (Vector Laboratories) and DAB as a chromagen. Brain sections and cultures were mounted in Mowiol-Dabco, organotypic slices in glycerol/PBS (2/1 vol/vol).

***In situ* Hybridization**

*In situ* Hybridization was performed on 150 μm-thick free-floating vibratome sections according to the protocol described in (Bellion et al., 2002) for whole brains.

**Microscopy.**

Brain sections and cultures were observed with a LEICA DM6000 upright fluorescent microscope. Grafted slices were imaged with a confocal (Leica TCS SP5) microscope, using a x10 objective.

In utero electroporated brain sections were imaged with an inverted microscope equipped with a spinning disk (Leica DMI4000) using a x20 long distance objective, or with a Leica TCS SP5 II confocal microscope, using a x40 objective. The image reconstruction of the brain half-hemisphere was performed respectively using the “photomerge” function of Adobe Photoshop or using the automatic “stitching” function of the Leica TCS SP5 II confocal microscope.

**Image processing.**

Cell counting in a volume or surface, area measurement, leading process orientation and length, cells trajectories and speed of migration in time-lapse experiments were performed with ImageJ (NIH, USA), especially with the Cell Counter, MtrackJ and NeuronJ plugins, or Metamorph software. Leading processes of eGFP+ MGE cells electroporated with eGFP and PAK3 constructs and grafted in cortical slices were traced on confocal pictures of the grafted cortical slices using the NeuronJ plugin of ImageJ. Leading process orientation was measured with regards to the cortical surface as explained in Baudoin et al (2012).

**Statistical analyses.**

In text and figures, the variability around mean values is represented by s.e.m. (standard error of mean). Statistical analysis were perfomed with GraphPad Prism or R software using Student’s t , Mann-Whitney,Chi2 tests, One or Two-way ANOVA followed by Post-hoc test (Dunn, Kruskal Wallis) and Poisson-anova model followed by likehood ratio test, across n individuals, where n is the number of embryos or cells as specified in the figure legend. Values of p<0.05 were considered significant and were indicated by a star within the area representing the corresponding value in cumulative histogram. In figures, levels of significance were expressed by * for p<0.05, ** for p<0.01, , *** for p<0.001 and **** for p<0.0001.
